# Homogeneity of antibody-drug conjugates critically impacts the therapeutic efficacy in brain tumors

**DOI:** 10.1101/2021.12.23.474044

**Authors:** Yasuaki Anami, Yoshihiro Otani, Wei Xiong, Summer Y. Y. Ha, Aiko Yamaguchi, Ningyan Zhang, Zhiqiang An, Balveen Kaur, Kyoji Tsuchikama

## Abstract

Glioblastoma multiforme (GBM) is characterized by aggressive growth and the poorest prognosis of all brain tumor types. Most therapies rarely provide clinically meaningful improvements in outcomes of patients with GBM. Antibody-drug conjugates (ADCs) are emerging chemotherapeutics with stunning success in cancer management. Although promising, clinical studies of three ADCs for treating GBM, including Depatux-M, have been discontinued because of safety concerns and limited therapeutic benefits. Here, we report that ADC homogeneity is a critical parameter to maximize the therapeutic potential in GBM therapy. We demonstrate that homogeneous conjugates generated using our linker show enhanced drug delivery to intracranial brain tumors. Notably, compared to heterogeneous ADCs, including a Depatux-M analog, our ADCs provide greatly improved antitumor effects and survival benefits in orthotopic brain tumor models, including a patient-derived xenograft model of GBM. Our findings warrant the future development of homogeneous ADCs as promising molecular entities toward cures for intractable brain tumors.

## INTRODUCTION

Glioblastoma multiforme (GBM) is the most aggressive brain tumor characterized by infiltrative growth to normal tissues, high proliferation rate, abundant angiogenesis, and intratumor and inter-patient heterogeneity (Inda et al., 2014; Parker et al., 2015; Shergalis et al., 2018). GBM has poorer survival rates than all other brain tumors (median survival time: 15-16 months) (Chinot et al., 2014; Stupp et al., 2005, 2009) due to quick relapse after standard therapy, namely surgical removal in combination with radiation therapy, chemotherapy using temozolomide, and/or tumor-treating fields. Deep infiltration of GBM into normal brain tissues makes complete surgical resection of tumor lesions a challenging task. While surgery is a proven option for primary GBM, its clinical benefit for patients with relapsed GBM remains unvalidated (Weller et al., 2014). To improve patients’ survival and quality of life, effective systemic therapies that can complement other treatment options are urgently needed.

Antibody-drug conjugates (ADCs) are an emerging class of chemotherapeutic agents consisting of tumor-targeting monoclonal antibodies (mAbs) with highly cytotoxic payloads attached through chemical linkers. ADCs can exert a durable and tumor-specific therapeutic effect by ensuring the delivery of conjugated cytotoxic payloads to antigen-positive tumor cells. Eleven ADCs have been approved by the U.S. Food and Drug Administration (FDA) (Dhillon, 2018; Drago et al., 2021; Mullard, 2021), and more than 100 ADCs are currently in clinical trials (Chau et al., 2019). Despite the success in the management of other cancers, ADCs have not yet shown remarkable treatment outcomes in patients with GBM. Three ADCs have advanced to clinical trials for GBM therapy: depatuxizumab mafodotin (Depatux-M or ABT-414) (Phillips et al., 2016), ABBV-221 (Phillips et al., 2018), and AMG-595 (Hamblett et al., 2015). These ADCs target EGFR and its active mutant EGFR variant III (EGFRvIII), which are signature receptors expressed in a subset of GBM tumors (Brennan et al., 2013). Unfortunately, clinical trials of the three ADCs have been terminated or discontinued (Newman, 2019; Rosenthal et al., 2019; Van Den Bent et al., 2020). No survival benefit was confirmed in a Phase 3 trial evaluating Depatux-M in patients with newly diagnosed GBM (Van Den Bent et al., 2020). In a preclinical study, ABBV-221 demonstrated greater treatment efficacy than could be achieved with Depatux-M; however, a Phase 1 study has raised safety concerns (Newman, 2019). The development of AMG-595 was discontinued upon completion of a Phase 1 study due to limited efficacy. Unlike other solid tumors, efficient mAb delivery to the brain is particularly challenging because of the blood–brain barrier (BBB), a tightly constituted endothelial cell border restricting the influx of large molecules from the vasculature to the brain parenchyma (Abbott et al., 2010; Banks, 2016). Therefore, to establish ADC-based GBM therapy as a practical clinical option, identifying and optimizing molecular parameters that negatively influence BBB permeability, therapeutic efficacy, and safety profiles are critically important.

Herein, we report that ADC homogeneity plays a critical role in payload delivery to intracranial brain tumors. We demonstrate that homogeneous ADCs elicit improved antitumor activity in intracranial brain tumor-bearing mouse models compared with heterogeneous variants prepared by stochastic cysteine– maleimide or lysine–amide coupling. We also show using mouse models how homogeneous conjugation at an optimal drug-to-antibody ratio (DAR) improves efficiency in payload delivery to intracranial GBM tumors, leading to dramatically extended survival. This finding suggests that ensuring ADC homogeneity is a crucial step to achieving clinically meaningful treatment outcomes in brain tumors, including GBM.

## RESULTS

### Construction of anti-EGFR ADCs with varied homogeneity

We have previously established click chemistry-empowered branched linkers for installing two identical or different payloads onto a single antibody in a site-specific and quantitative manner (Anami and Tsuchikama, 2020; Anami et al., 2017; Yamazaki et al., 2021). We have also developed the glutamic acid–valine–citrulline (EVCit) cleavable linker enabling the intracellular release of payloads in a traceless fashion while minimizing premature linker degradation in human and mouse plasma (Anami et al., 2018). Indeed, we have confirmed that the maximum tolerated dose of an EVCit-based dual-drug ADC containing monomethyl auristatin E (MMAE) and monomethyl auristatin F (MMAF) is higher than 40 mg/kg in non-tumor bearing mice (Yamazaki et al., 2021). Using these technologies, we set out to construct a homogeneous ADC targeting both EGFR and EGFRvIII (**Figure 1A**). We used cetuximab with N88A and N297A double mutations for ADC construction. Cetuximab, a human–murine chimeric mAb targeting the extracellular domain III of EGFR and EGFRvIII, has been approved for the treatment of colorectal cancer and head and neck cancer (EliLilly, 2004). The N88A/N297A double mutations remove two *N*-glycans on the side chains of asparagine 88 within the Fab moiety and asparagine 297 within the Fc moiety (Giddens et al., 2018). Thus, this modification allows the omission of the deglycosylation step required for following microbial transglutaminase (MTGase)-mediated linker conjugation. In addition, *N*-glycan removal abrogates immune responses derived from interactions with Fcγ receptors expressed in immune cells, which can minimize undesired systemic toxicity or inflammatory response (Herbst et al., 2020; White et al., 2020). Atezolizumab (TECENTRIQ^®^, anti-PD-L1 mAb) is a recent example with an N297A mutation approved by the FDA.

**Figure 1.**
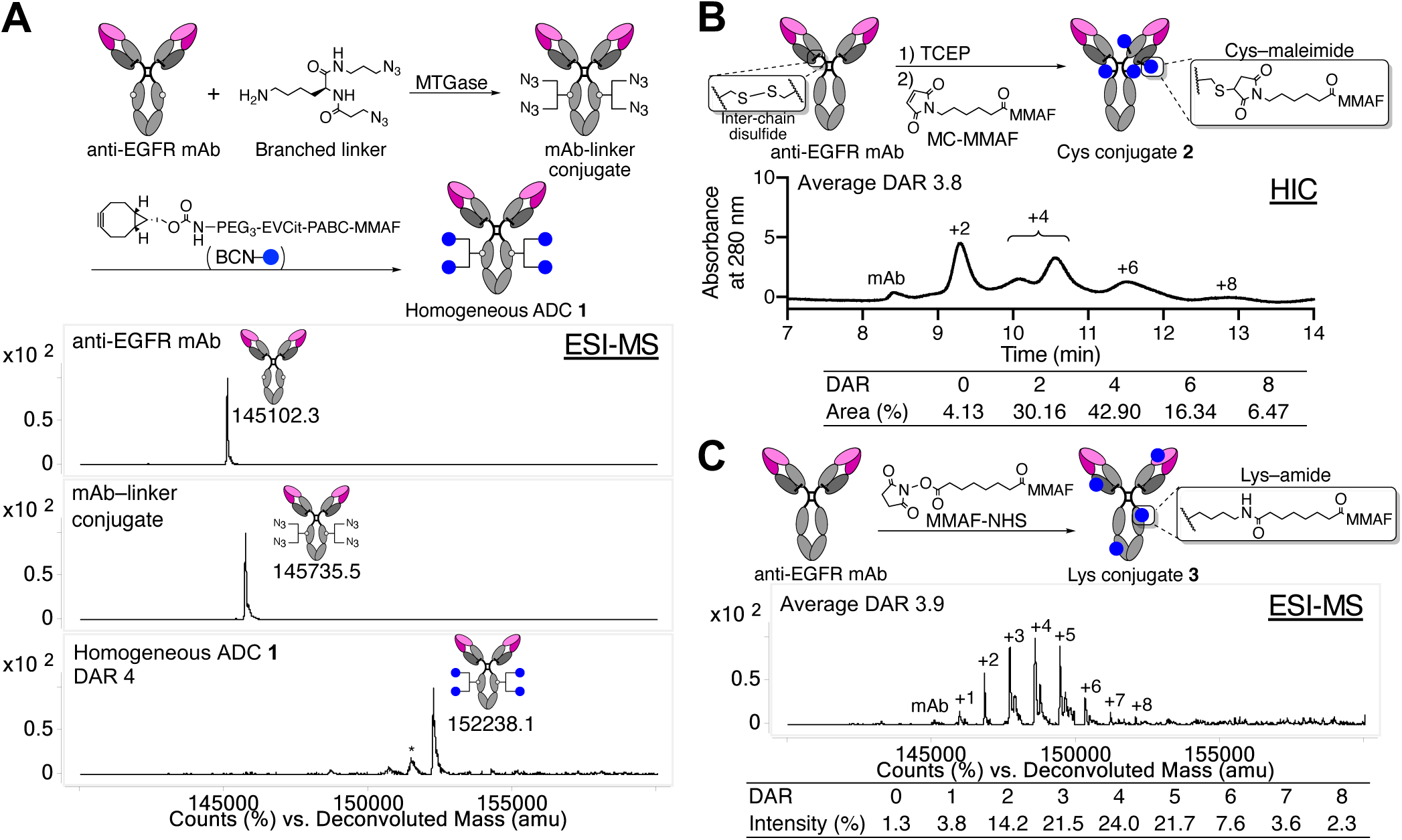
Construction and characterization of anti-EGFR ADCs. **A** Preparation and ESI-MS analysis of homogeneous ADC **1**. Top panel: N88A/N297A anti-EGFR mAb (cetuximab mutant). Middle panel: mAb–linker conjugate. Bottom panel: homogeneous ADC **1** with a DAR of 4. Asterisk (*) indicates a fragment ion detected in ESI-MS analysis. **B** Preparation and HIC analysis of Cys conjugate **2** under physiological conditions (phosphate buffer, pH 7.4). The average DAR was determined to be 3.8 based on UV peak area of each DAR species. **C** Preparation and ESI-MS analysis of Lys conjugate **3**. The average DAR was determined to be 3.9 based on the ion intensity of each DAR species. BCN, bicyclo[6.1.0]nonyne; DAR, drug-to-antibody ratio; ESI-MS, electrospray ionization mass spectrometry; MC, maleimidecaproyl; MMAF, monomethyl auristatin F; MTGase, microbial transglutaminase; HIC, hydrophobicity interaction chromatography; NHS, *N*-hydroxysuccinimide; PABC, *p*-aminobenzyloxycarbonyl; PEG, polyethylene glycol; TCEP, tris(2-carboxyethyl)phosphine.

We began the ADC construction by installing branched diazide linkers site-specifically onto glutamine 295 (Q295) within the parent N88A/N297A anti-EGFR mAb using MTGase (Anami and Tsuchikama, 2020) (**Figure 1A**). This enzymatic conjugation yielded a homogeneous mAb–branched linker conjugate in high yield. In parallel, we synthesized a payload module consisting of bicyclo[6.1.0]nonyne (BCN, as a reaction handle for following strain-promoted azide–alkyne click reaction), EVCit (as a cathepsin-responsive cleavable sequence), *p*-aminobenzyloxycarbonyl (PABC) spacer, and MMAF (BCN–EVCit–PABC–MMAF, see Supplementary Information for synthesis details). Finally, the click reaction between the azide groups on the branched linkers and BCN–EVCit–PABC– MMAF quantitatively afforded anti-EGFR ADC **1** with a DAR of 4. We confirmed the homogeneity of ADC **1** by reverse-phase HPLC and electrospray ionization mass spectrometry (ESI-MS) analysis (**Figure S1A**). Using the same parent anti-EGFR mAb, we also prepared two heterogeneous variants that resemble the structure of Depatux-M (Cys conjugate **2**) (Phillips et al., 2016) or the conjugation modality of AMG-595 (Lys conjugate **3**) (Hamblett et al., 2015). For the preparation of Cys conjugate **2**, non-cleavable maleimidocaproyl MMAF (MC–MMAF) was installed by partial disulfide bond reduction and following cysteine–maleimide alkylation. We confirmed by hydrophobic interaction chromatography (HIC) analysis that Cys conjugate **2** consisted of DAR-0, 2, 4, 6, and 8 species (average DAR: 3.8, **Figure 1B**). To prepare Lys conjugate **3**, we synthesized and used non-cleavable MMAF–*N*-hydroxysuccinimide (NHS) ester for lysine coupling-based conjugation (See Supplementary Information for synthesis details). ESI-MS analysis revealed that this heterogeneous conjugate consisted of multiple products with DARs ranging from 0 to 8 (average DAR: 3.9, **Figure 1C**).

### Cysteine–maleimide conjugation does not impair EGFR-specific potency *in vitro* but reduces long-term stability

Size-exclusion chromatography (SEC) analysis revealed that all ADCs generated predominantly existed in the monomer form (**Figure S1B**). These ADCs were also tested for long-term stability under physiological conditions by being incubated at 37 °C in PBS (pH 7.4) for 28 days. We observed no significant degradation or aggregation for homogeneous ADC **1** and Lys conjugate **3** (**Figure S2**). In contrast, Cys conjugate **2** showed two new peaks after the peak corresponding to its monomeric form, indicating that fragmentation or partial dissociation of the heavy and light chains occurred. These results suggest that both MTGase-mediated homogeneous conjugation and lysine coupling offer higher thermal stability compared to the one achieved by cysteine–maleimide conjugation.

Next, we assessed antigen-specific binding of the ADCs by cell-based ELISA (**Figure 2A** and Table **S1**). All ADCs showed binding affinities for EGFRvIII-positive U87ΔEGFR-luc cells (K_D_: 0.044–0.047 nM) comparable to that of the unmodified N88A/N297A cetuximab (K_D_: 0.039 nM). In addition, none of the ADCs bound to EGFR-negative HEK293 cells. These results demonstrate that the ADCs retained their binding affinity and specificity regardless of conjugation methods. We also tested these conjugates for cell killing potency in U87ΔEGFR-luc, Gli36δEGFR (EGFRvIII-positive), and HEK293 cells (EGFR-negative control) (**Figure 2B** and **Table S2**). All DAR 4 ADCs showed comparable potency in the EGFRvIII-positive GBM cells (EC_50_ values: 0.072–0.140 nM in U87ΔEGFR-luc and 0.035–0.048 nM in Gli36δEGFR cells), but not in HEK293 cells. This result is in line with previous reports demonstrating that MMAF ADCs can exert pM-level cell killing potency with or without a cleavable linker (Deonarain et al., 2014; Doronina et al., 2006).

**Figure 2.**
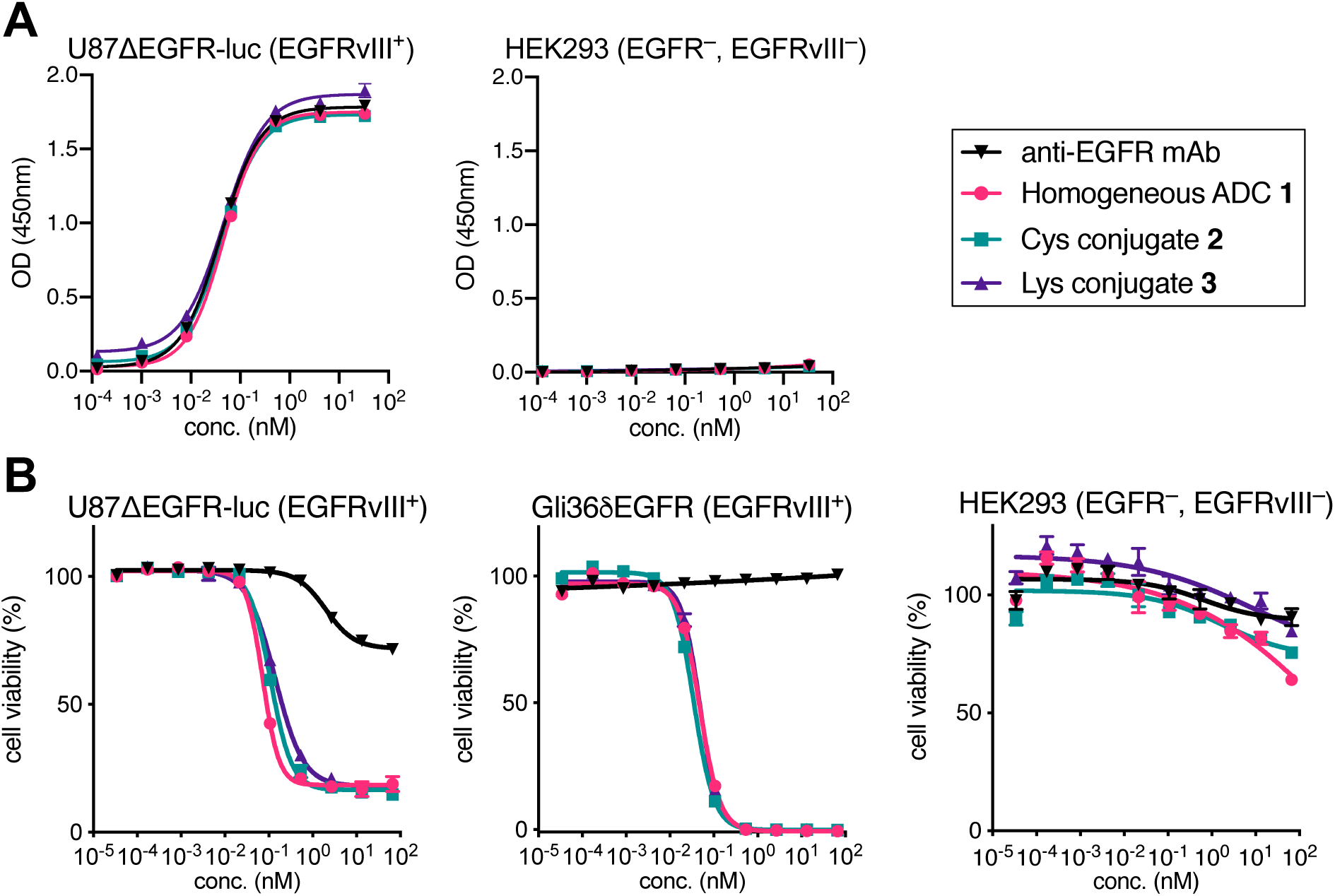
Evaluation of antigen-specific binding and *in vitro* cytotoxicity. **A** Cell-based ELISA in U87ΔEGFR-luc (EGFRvIII^+^) and HEK293 (wtEGFR^−^, EGFRvIII^−^) cells. **B** Cell killing potency in U87ΔEGFR-luc, Gli36δEGFR (EGFRvIII^+^), and HEK293. We tested the parent anti-EGFR mAb (black inversed triangle), homogeneous ADC **1** (magenta circle), Cys conjugate **2** (green square), and Lys conjugate **3** (purple triangle). Concentrations are based on the antibody dose without normalizing to each DAR. All assays were performed in triplicate. Data are presented as mean values ± SEM.

### The homogeneous anti-EGFR ADC exerts significantly improved therapeutic efficacy in orthotopic mouse models of GBM

To evaluate the *in vivo* anti-tumor activity of the three anti-EGFR ADCs, we first performed a treatment study using a cell line-derived xenograft model of human GBM. To gain clinically translatable insights into the influence of conjugation modality on drug delivery to brain tumors, intracranially implanted models were used instead of subcutaneous models. NOD*scid* gamma (NSG) mice were orthotopically implanted with U87ΔEGFR-luc cells and injected intravenously with a single dose of each ADC (homogeneous ADC **1**, Cys conjugate **2**, or Lys conjugate **3**) at 3 mg/kg 5 days post-implantation (**Figure 3A**). Tumor growth and body weight were monitored periodically (**Figure S3**). No acute toxicity associated with ADC administration was observed in either group over the course of the study (**Figure S3A**). The short survival time observed for the untreated group (median survival: 14 days, **Figure 3B**) demonstrates the extremely aggressive growth of this GBM model. Homogeneous ADC **1** exerted remarkable antitumor activity with statistically significant survival benefits; the median survival rate increased from 14 days in the untreated group to 43 days in the treated cohort (207% extension, *P* = 0.0032). In addition, two out of six mice survived at the end of the study (Day 60) with no detectable bioluminescence signal from implanted tumors (**Figure S3B**), indicating that these two mice achieved complete remission. In contrast, the heterogeneous conjugates exhibited limited therapeutic effects with marginally increased median survival times (22 days, Cys conjugate **2** and 23.5 days, Lys conjugate **3**), which were inferior to that provided by homogeneous ADC **1** (*P* = 0.0046). Indeed, all mice in these two groups died or reached the pre-defined humane endpoint by the end of the study (**Figure 3B**). This result is in contrast to our observation that ADCs **1**–**3** showed comparable *in vitro* cell killing potency in U87ΔEGFR-luc cells (**Figure 2B**).

**Figure 3.**
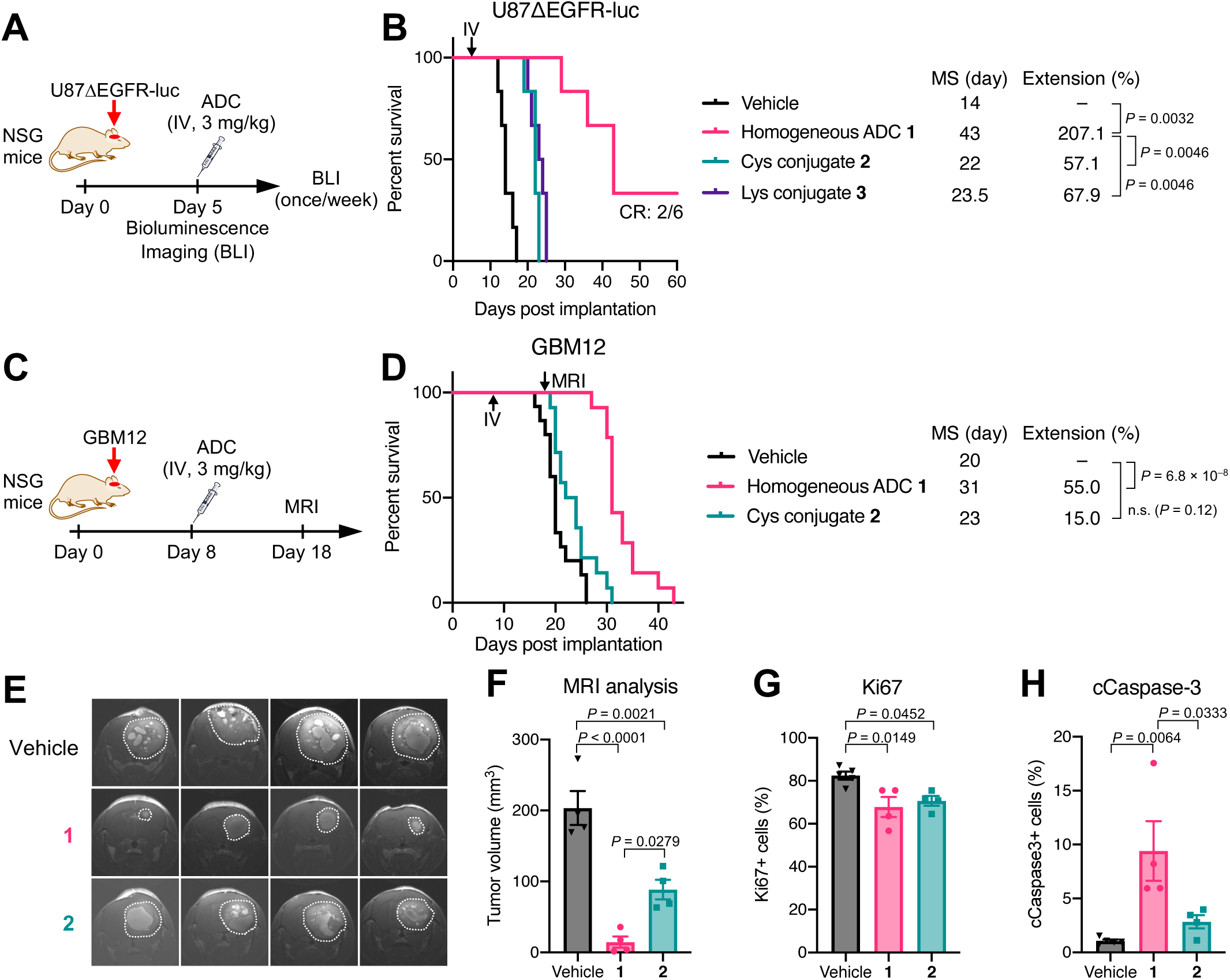
ADC homogeneity enhances therapeutic efficacy in orthotopic GBM mouse models. **A** Study schedule in the U87ΔEGFR-luc xenograft model. Male and female NSG mice were intracranially implanted with U87ΔEGFR-luc cells. Five days after tumor implantation, each group was treated with a single intravenous administration of each ADC at 3 mg/kg and monitored by BLI once a week. **B** Survival curves in the U87ΔEGFR-luc model (n = 6/group). Vehicle (black), homogeneous ADC **1** (magenta), Cys conjugate **2** (green), and Lys conjugate **3** (purple). Two out of six mice treated with homogeneous ADC **1** survived over 60 days without detectable tumor-derived bioluminescence signal. All animals other than the ones that were found dead or achieved complete remission were euthanized at the pre-defined humane endpoint, which were counted as deaths. **C** Study schedule in the GBM12 PDX model. Tumor-bearing NSG mice were treated with ADC **1** or **2** at 3 mg/kg 8 days post-implantation. MRI was performed on Day 18. **D** Survival curves in the GBM12 model (n = 15 for vehicle; n = 14 for ADCs). **E** Coronal MRI images on Day 18 (n = 4). Tumor lesions are indicated with white dots. **F** Estimated tumor volume by MRI image-based quantification (n = 4). **G** Population of Ki67-positive cells in the GBM12 tumors harvested at the terminal stage (n = 5 for vehicle, n = 4 for ADCs). **H** Population of cCaspase-3-positive cells in the GBM12 tumors harvested at the terminal stage (n = 5 for vehicle, n = 4 for ADCs). Data are presented as mean values ± SEM. Kaplan-Meier survival curve statistics were analyzed with a log-rank test. To control the family-wise error rate in multiple comparisons, crude *P* values were adjusted by the Bonferroni method. For MRI and tissue analysis, a one-way ANOVA with a Tukey-Kramer post hoc test was used (see **Table S4** for details). BLI, bioluminescence imaging; cCaspase-3, cleaved caspase-3; CR, complete remission; MRI, magnetic resonance imaging; MS, median survival; PDX, patient-derived xenograft.

To further validate the therapeutic potential of homogeneous ADC **1**, we sought to use a patient-derived xenograft (PDX) tumor model of GBM. PDX models maintain pathohistological and genetic properties of original tumors as well as therapeutic responses to anti-cancer treatments. As such, PDX models provide clinically relevant and translatable data (Hidalgo et al., 2014). To this end, we used GBM12, a PDX model of GBM overexpressing wild-type EGFR (Sarkaria et al., 2006). A study has shown that GBM12 tumors show heterogeneous BBB disruption, meaning that some GBM12 tumor cells are likely protected by an intact BBB (Parrish et al., 2015). Before initiating an *in vivo* assessment, homogeneous ADC **1** and heterogeneous Cys conjugate **2** were evaluated for cell killing potency in GBM12 cells. Both ADCs efficiently killed GBM12 cells with comparable EC_50_ values (homogeneous ADC **1**: 0.08 nM, Cys conjugate **2**: 0.11 nM, **Figure S4A**). Next, we investigated whether or not homogeneous conjugate **1** also showed a greater treatment effect in the orthotopic GBM12 mouse model than could be achieved by Cys conjugate **2**. NSG mice bearing intracranial GBM12 tumors were injected intravenously with a single dose of either conjugate (3 mg/kg) 8 days post-implantation (**Figure 3C**). Tumor size was noninvasively measured by magnetic resonance imaging (MRI) on Day 18. No acute toxicity was observed in either group over the course of the study (**Figure S4B**). Homogeneous ADC **1** effectively suppressed tumor growth with a statistically significant survival benefit (median survival: 31 days, +55% relative to the vehicle group, *P* = 6.8 × 10^−8^), whereas Cys conjugate **2** showed a marginal therapeutic effect (median survival: 23 days, +15% extension, *P* = 0.12, **Figure 3D**). MRI on Day 18 showed that the tumors treated with homogeneous ADC **1** (average size: 14.71 ± 7.90 mm^3^) were markedly smaller than the untreated ones (average size: 203.46 ± 23.81 mm^3^, *P* < 0.0001, **Figures 3E** and **3F**). Cys conjugate **2** also inhibited tumor growth (average size: 88.63 ± 13.71 mm^3^) but less effectively than homogeneous ADC **1** (*P* = 0.0279). To investigate how each ADC influenced cell proliferation and apoptosis, we performed immunohistochemistry analysis of brain tissues harvested from each group at the terminal stage (vehicle: 20–26 days, ADC **1**: 30–35 days; Cys conjugate **2**: 24–31 days, **Figures S4C–F**). About 80% of cells were Ki67-positive in the vehicle-treated group, while about 70% of cells were Ki67-positive in both ADC-treated groups (**Figure 3G**). This result indicates that antiproliferative effects by both ADCs declined to similar levels at the terminal stage. In contrast, the population of cleaved caspase-3 (cCaspase-3)-positive cells in the tumors treated with ADC **1** (9.4 ± 2.8%) was significantly higher than that in the tumors treated with vehicle (1.1 ± 0.1%, *P* = 0.0064) or Cys conjugate **2** (2.8 ± 0.6%, *P* = 0.0333, **Figure 3H**), suggesting that homogeneous ADC **1** induced apoptosis more effectively than heterogeneous ADC **2** over the course of the study. Given that the histopathology analysis was performed at the terminal stage of each group, more significant differences in Ki67 and cCaspase-3 levels could have been observed at the same time point in the early stage. Collectively, these results demonstrate that homogeneous ADC **1** can eradicate intracranial GBM tumors more efficiently than its heterogeneous variants.

### Homogeneity also improves *in vivo* therapeutic efficacy of other ADCs for EGFRvIII- and HER2-positive brain tumors

To generalize our findings, we tested other homogeneous ADCs for treatment efficacy in orthotopic brain tumor models. To this end, homogeneous anti-EGFRvIII ADC **4** (DAR 4) and heterogeneous variant **5** (Lys conjugate, average DAR: 4.7) were constructed from N297A depatuxizumab, the parent mAb of Depatux-M (Phillips et al., 2016) (**Figure S5A**). Both ADCs showed comparable cell killing potency in EGFRvIII-positive U87ΔEGFR-luc cells (EC_50_ values: 0.15 nM, homogeneous ADC **4** and 0.26 nM, Lys conjugate **5, Figure S5B**). Subsequently, NSG mice bearing intracranial U87ΔEGFR-luc tumors were treated with a single dose of each ADC at 3 mg/kg 5 days after tumor implantation (**Figure 4A**). Homogeneous ADC **4** showed a remarkable survival benefit (median survival: >60 days). Four out of six mice treated with homogeneous ADC **4** survived over the course of the study. In addition, MRI on Day 61 showed no detectable brain tumor lesion in these survivors, indicating complete remission (**Figure S5C**). In contrast, heterogeneous variant **5** extended median survival time (33 days) less significantly than homogeneous ADC **4** (*P* = 0.0195, **Figure 4B**).

**Figure 4.**
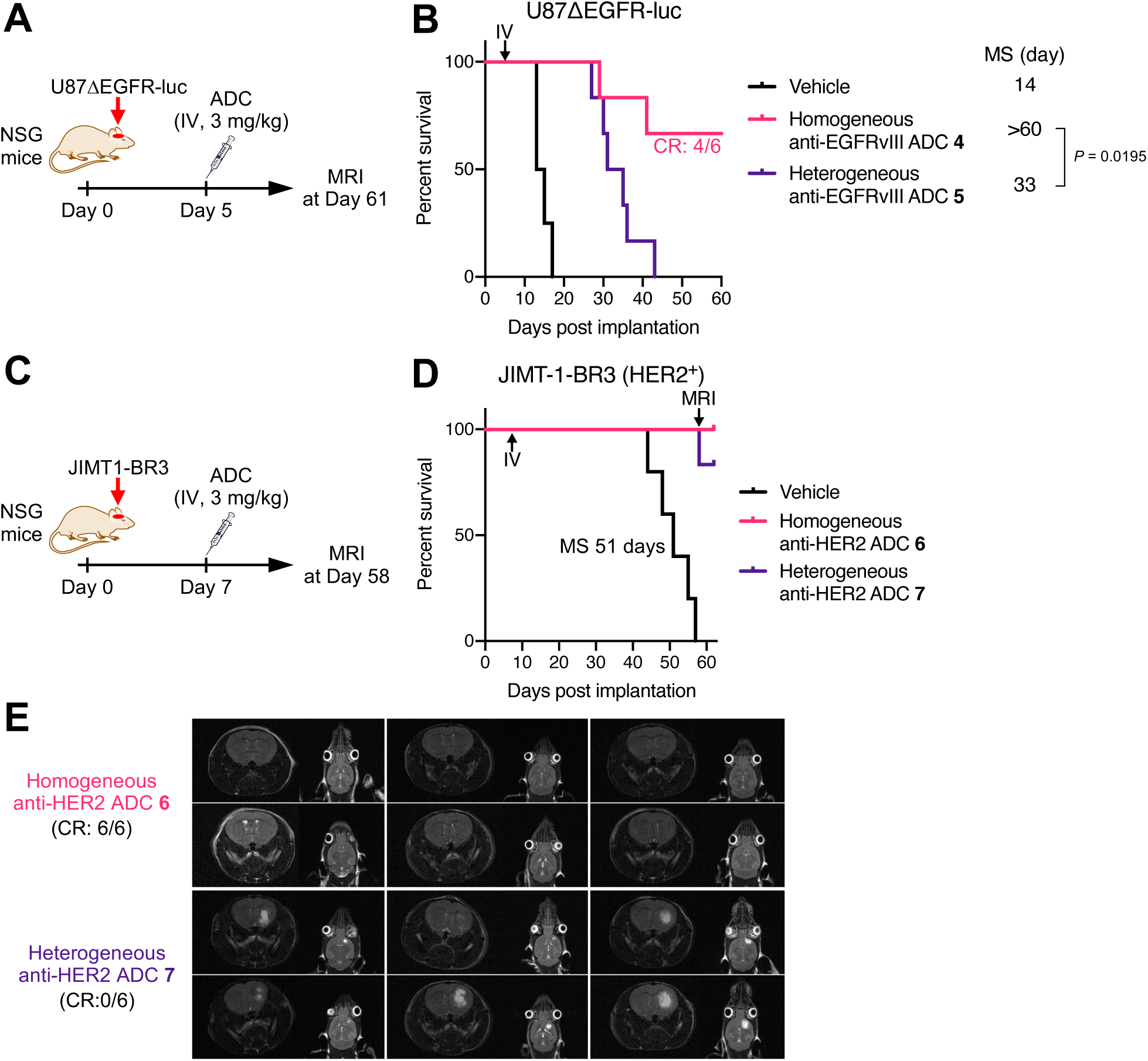
Homogeneous depatuxizumab- and trastuzumab-based ADCs show enhanced therapeutic efficacy in orthotopic brain tumor mouse models. **A** Study schedule in the orthotopic U87ΔEGFR-luc xenograft mouse model. Male and female NSG mice were intracranially implanted with U87ΔEGFR-luc cells. Five days after tumor implantation, each group was treated with a single intravenous administration of each depatuxizumab-based anti-EGFRvIII ADC at 3 mg/kg. MRI was performed on Day 61. **B** Survival curves in the U87ΔEGFR-luc model. Vehicle (black, n = 4), homogeneous ADC **4** (magenta, n = 6), and heterogeneous Lys conjugate **5** (purple, n = 6). Four out of six mice treated with homogeneous ADC **4** survived over 60 days without detectable tumor lesions (**Figure S5C**). All animals other than the ones that were found dead or achieved complete remission were euthanized at the pre-defined humane endpoint, which were counted as deaths. **C** Study schedule for the intracranially implanted JIMT1-BR3 tumor mouse model. NSG mice bearing intracranial JIMT-1-BR3 tumors were injected intravenously with a single dose of each trastuzumab-based anti-HER2 ADC at 3 mg/kg 7 days post-implantation. MRI was performed on Day 58. **D** Survival curves in the JIMT-1-BR3 model (n = 6). Vehicle (black), homogeneous anti-HER2 ADC **6** (magenta), and heterogeneous Lys conjugate **7** (purple). **E** Coronal and sagittal MRI images of the intracranial JIMT-1-BR3 tumor-bearing mice on Day 58. Brain tumor lesions were detected in the mice treated with heterogeneous ADC **7** (CR: 0/6) but not in those treated with homogeneous ADC **6** (CR: 6/6). Kaplan-Meier survival curve statistics were analyzed with a log-rank test (see **Table S4** for details).

Next, we performed similar *in vitro* and *in vivo* studies using a HER2-positive brain tumor model. Brain metastasis is observed in 25–50% patients with advanced HER2-positive breast tumors (Zimmer et al., 2020), representing a difficult-to-treat population. We prepared anti-HER2 homogeneous ADC **6** (DAR 4) and a heterogeneous variant (Lys conjugate **7**, average DAR: 4.2) from N297A trastuzumab and evaluated their cell killing potency in HER2-positive JIMT-1-BR3 cells (**Figures S6A** and **S6B**). JIMT-1-BR3 is a HER2-positive breast cancer cell line established from a subpopulation of the parent JIMT-1 cells that metastasized to the brain in a rodent model (Palmieri et al., 2014). Both ADCs efficiently killed JIMT-1-BR3 cells with comparable EC_50_ values (homogeneous ADC **6**: 0.037 nM, Lys-conjugate **7**: 0.059 nM, **Figure S6B**). Subsequently, NSG mice bearing intracranial JIMT-1-BR3 tumors were injected intravenously with each ADC at 3 mg/kg 7 days post tumor implantation (**Figure 4C**). Most mice in both ADC groups survived over the course of the study (Homogeneous ADC **4**: all mice, heterogeneous ADC **5**: 5 out of 6 mice), while the median survival time without treatment was 51 days (**Figure 4D**). However, MRI on Day 58 revealed a clear difference in efficacy between these two ADC groups; tumor lesions were detected in all mice treated with heterogeneous ADC **7** but not in those treated with homogeneous ADC **6** (**Figure 4E**). Hematoxylin and eosin (H&E) staining of the brain tissues also validated this observation (**Figure S6C**). Collectively, these findings strongly support our hypothesis that the use of homogeneous ADCs can lead to significantly improved treatment outcomes in a broad range of brain tumors compared to heterogeneous ADC-based treatment.

### Clearance and linker stability in circulation are not the primary factors reducing the efficiency in payload delivery to brain tumors

To understand how the antibody–drug conjugation modality impacts overall efficacy in the treatment of brain tumors, we set out to assess *in vivo* pharmacokinetic (PK) profiles of selected ADCs. Homogeneous anti-EGFR ADC **1**, heterogeneous Cys conjugate **2**, heterogeneous Lys conjugate **3**, or the parental anti-EGFR mAb (3 mg/kg) was intravenously administered into CD-1 mice^®^. Subsequently, blood samples were periodically collected. The concentrations of both total mAb and intact ADC were then determined by sandwich ELISA. In total mAb analysis, homogeneous ADC **1** and Lys conjugate **3** showed half-lives at the elimination phase (t_1/2β_ = 9.8 days, ADC **1** and t_1/2β_ = 10.4 days, Lys conjugate **3**) comparable with that of the unmodified mAb (10.9 days), whereas Cys conjugate **2** showed a slightly decreased half-life (t_1/2β_ = 7.8 days, **Figure 5A** and **Table S3**). We found that Cys conjugate **2** showed thermal instability after a 28-day incubation under physiological conditions probably due to partly cleaved interchain disulfide bonds (**Figure S2C**). This instability may account for the increased clearance rate. In intact ADC analysis, no significant decrease in half-lives was observed for homogeneous ADC **1** (t_1/2β_ = 8.6 days) or Lys conjugate **3** (t_1/2β_ = 8.9 days), indicating that there was almost no premature release of MMAF during circulation (**Figure 5B** and **Table S3**). In contrast, the concentration of intact Cys conjugate **2** declined at a faster rate (t_1/2β_ = 4.2 days), indicating that the conjugated MMAF was partly lost in circulation. Previous reports have shown that cysteine-containing serum proteins such as albumin promote dissociation of cysteine–maleimide linkage within ADCs through a thiol exchange reaction, leading to partial deconjugation of payloads in circulation (Lyon et al., 2014; Tumey et al., 2014).

**Figure 5.**
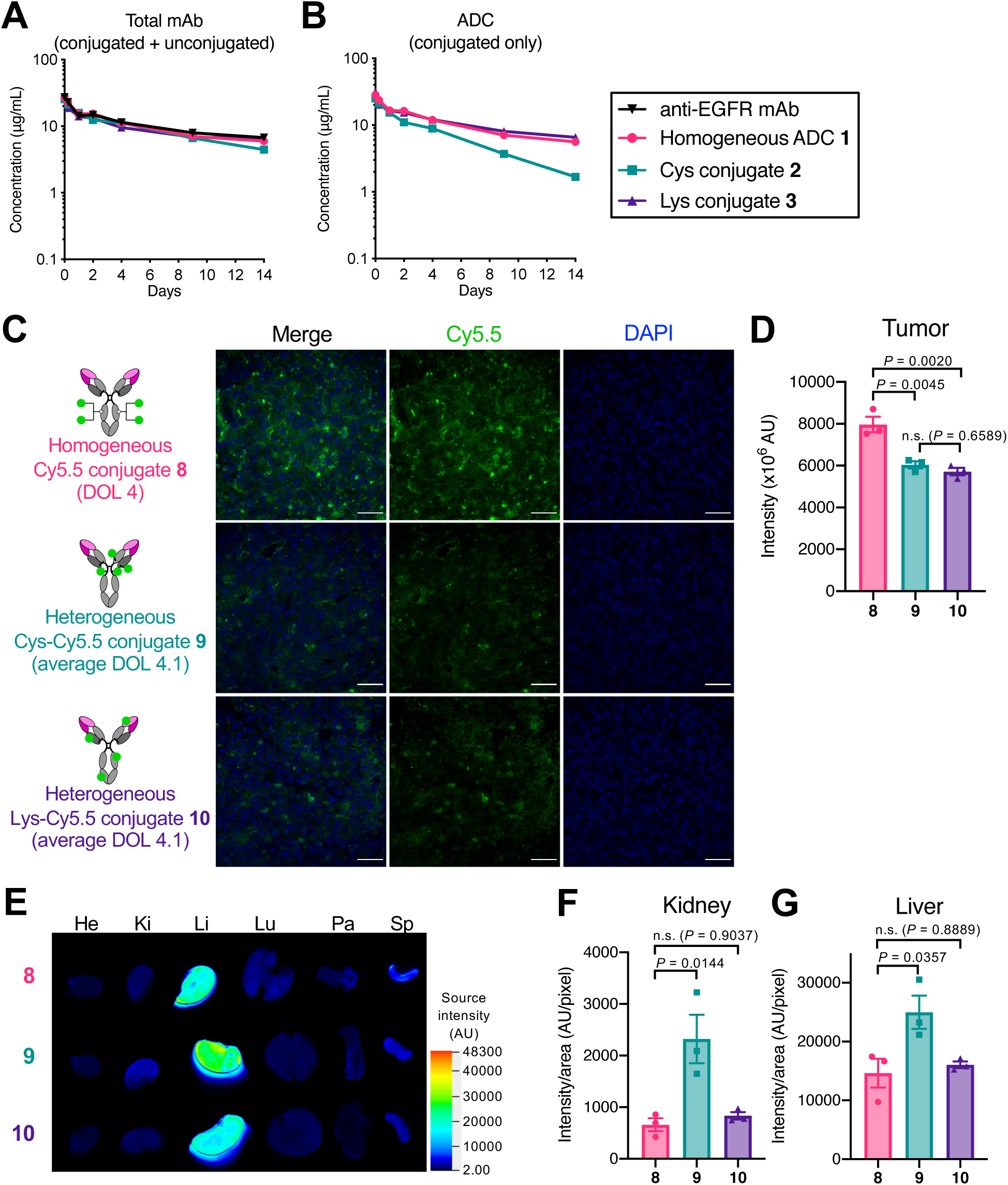
Promoted clearance and premature payload loss in circulation are not the primary factors attenuating the brain tumor-targeting efficiency of ADCs. **A**,**B** PK of unmodified N88A/N297A anti-EGFR mAb (black inversed triangle), homogeneous ADC **1** (magenta circle), Cys conjugate **2** (green square), and Lys conjugate **3** (purple triangle) in female CD-1^®^ mice (n = 3). At the indicated time points, blood was collected to quantify total antibody (conjugated and unconjugated, **A**) and intact ADC (conjugated only, **B**) by sandwich ELISA. **C** Fluorescence images of brain tumor tissues harvested 48 hours after injecting each Cy5.5 conjugate (n = 3, scale bar: 50 μm). **D** Semi-quantification of the Cy5.5 signal detected in the brain tumor tissues. Three regions of interest (ROI) were randomly selected in each tissue sample to calculate signal intensity. **E** Ex vivo fluorescence images of the other organs (He, heart; Ki, kidney; Li, liver; Lu, lung; Pa, pancreas; Sp, spleen) detected using a 700 nm channel (n = 3). **F**,**G** Semi-quantification of the Cy5.5 signal detected in the kidneys and liver. A representative image from each group is shown in all panels of fluorescence images. Data are presented as mean values ± SEM. For statistical analysis, a one-way ANOVA with a Tukey-Kramer post hoc test was used (see **Table S4** for details). DOL, degree of labeling.

As demonstrated above, promoted clearance and payload deconjugation may partly account for the poor treatment efficacy observed for Cys conjugate **2** in the orthotopic GBM models. However, these factors are likely irrelevant to the inferior efficacy observed for the lysine conjugates, which were designed not to show thermal instability or undergo deconjugation in circulation. To uncover other contributing factors, we performed a biodistribution study using the orthotopic U87ΔEGFR-luc xenograft mouse model. As surrogates of the ADCs we used earlier, the following fluorescent dye conjugates were prepared from the N88A/N297A cetuximab: homogeneous Cy5.5 conjugate **8** (degree of labeling or DOL: 4) and two heterogeneous Cy5.5 conjugates by cysteine–maleimide coupling (Cys-Cy5.5 conjugate **9**, average DOL: 4.1) and lysine coupling (Lys-Cy5.5 conjugate **10**, average DOL: 4.1, **Figures S7A–C**). We synthesized and used DBCO–EVCit–Cy5.5 module to construct homogeneous conjugate **8** (see Supplementary Information for synthesis details). In all cases, Cy5.5 was incorporated as a payload surrogate into the parent mAb with the same linkers and conjugation chemistries that were used to prepare corresponding ADCs. Orthotopic U87ΔEGFR-luc tumor-bearing NSG mice were administered intravenously with each dye conjugate at 3 mg/kg 5 days post tumor implantation. Two days after administration, all animals underwent cardiac perfusion for removing conjugates circulating or bound to the vascular endothelial cells. We then harvested major organs including tumor-bearing brains. Fluorescence imaging revealed that homogeneous Cy5.5 conjugate **8** accumulated in the brain tumors more effectively than heterogeneous conjugates **9** (*P* = 0.0045) and **10** (*P* = 0.0020, **Figures 5C** and **5D**). We did not see a significant difference in intracranial U87ΔEGFR-luc tumor targeting ability between Cys conjugate **9** and Lys conjugate **10**. We also confirmed in a separate biodistribution study that the cathepsin-responsive cleavage EVCit linker did not significantly contribute to the enhanced accumulation in U87ΔEGFR-luc tumors (**Figures S7D and S7E**). Cys-Cy5.5 conjugate **9** showed an increased fluorescent signal in the kidneys and liver compared to homogeneous Cy5.5 conjugate **8** (kidneys: *P* = 0.0144, liver: *P* = 0.0357), due probably to partial deconjugation of the maleimide-Cy5.5 modules in circulation and following hepatic and renal clearance (**Figures 5E–G**). However, we did not observe such increased liver and kidney accumulation for Lys-Cy5.5 conjugate **10**. Taken together, these findings suggest that promoted clearance of conjugated payloads and linker instability are not the primary factors attenuating the brain tumor-targeting efficiency.

### Homogeneous conjugation enables efficient payload delivery to intracranial tumors for days

We performed longitudinal intravital imaging to clarify spatiotemporal changes in the accumulation of payloads in brain tumors. GBM12 cells that stably express Red Fluorescent Protein (GBM12-RFP) were implanted into NSG mice intracranially, and either Cy5.5 conjugate **8, 9**, or **10** was administered intravenously at 3 mg/kg 14 days post-implantation (**Figure 6A**). Fluorescence images were then taken at multiple time points through a cranial window. As demonstrated by the increasing RFP signals, the implanted tumors continued to grow throughout the study (**Figures 6B–D**). To offset the intragroup and intergroup variances derived from tumor growth, the Cy5.5 signal intensity was normalized to the RFP signal intensity at each time point. The intratumor concentrations of the three conjugates peaked around Day 3 post-administration and then declined over time (**Figure 6E**). Notably, homogeneous conjugate **8** accumulated in the orthotopic GBM12 tumors more significantly and persistently than heterogeneous variants **9** and **10**; the statistically significant enhancement was observed for up to 5 days (**Figure 6E**). Overall, these results suggest that homogeneous conjugation allows intravenously administered antibody conjugates to target brain tumors with enhanced payload delivery efficiency and durability.

**Figure 6.**
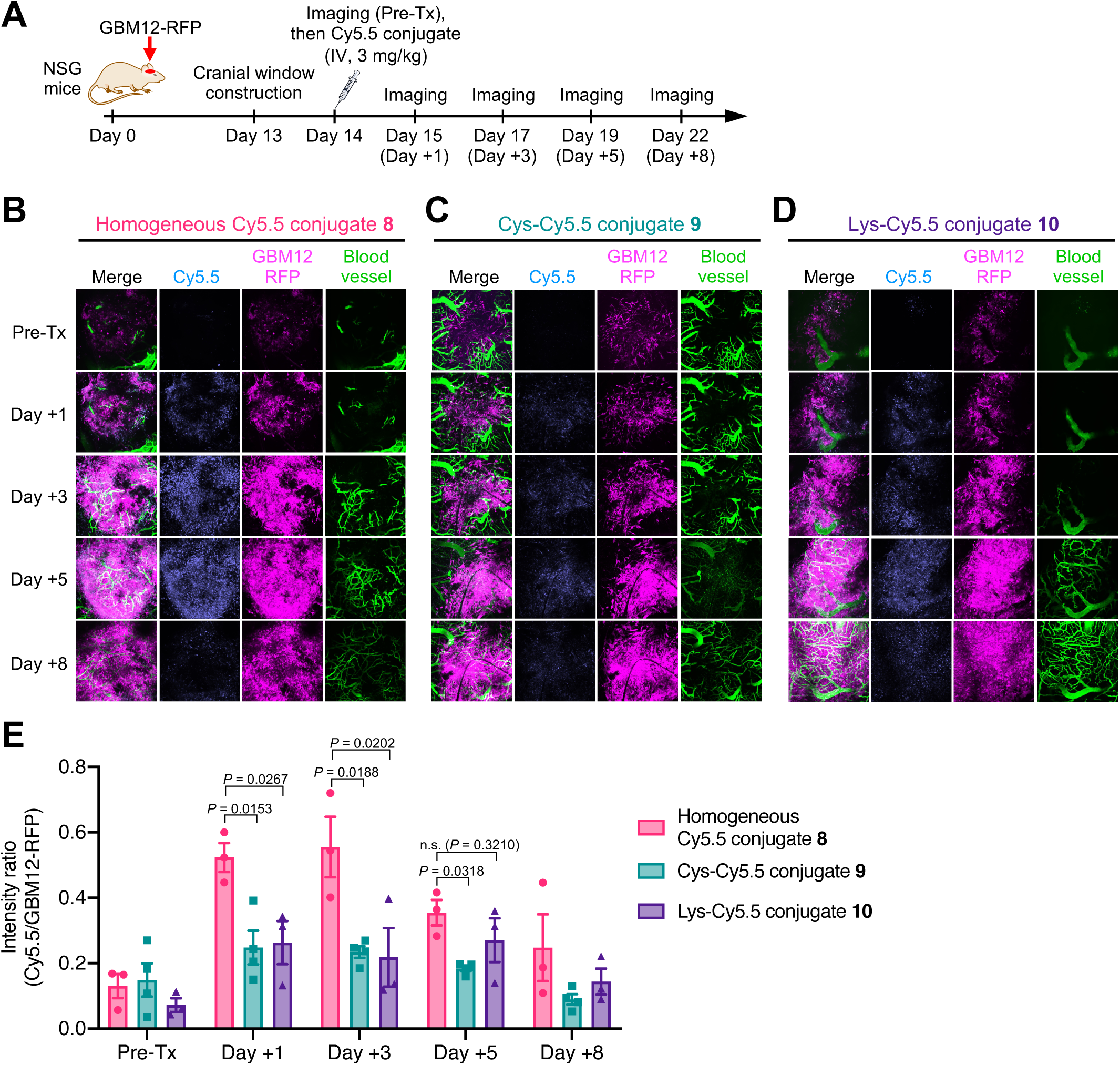
Homogeneous conjugation allows for enhanced payload delivery to orthotopically xenografted GBM tumors for several days. **A** Study schedule for intravital imaging. Male NSG mice were intracranially implanted with GBM12-RFP cells. Thirteen days after tumor implantation, a cranial window was constructed in each animal. Next day, each group was then injected intravenously with a single dose of each Cy5.5 conjugate at 3 mg/kg. Fluorescence images were taken before administration (Pre-Tx) and on Day 1, 3, 5, and 8 post-administration. FITC-conjugated dextran was injected right before each imaging session to visualize the brain microvasculature. **B** Intravital images of GBM12-RFP tumors treated with homogeneous Cy5.5 conjugate **8** (n = 3). A representative image at each time point is shown. **C** Intravital images of GBM12-RFP tumors treated with Cys-Cy5.5 conjugate **9** (n = 4). A representative image at each time point is shown. **D** Intravital images of GBM12-RFP tumors treated with Lys-Cy5.5 conjugate **10** (n = 3). A representative image at each time point is shown. Cy5.5, cyan; RFP, magenta; FITC, green. **E** Normalized Cy5.5 intensity (Cy5.5 signal/GBM12-RFP signal). Data are presented as mean values ± SEM. For statistical analysis, a one-way ANOVA with a Dunnett’s post hoc test (control: homogeneous conjugate **8**) was used (see **Table S4** for details).

### High-DAR components in heterogeneous MMAF ADCs target brain tumors less efficiently than low-DAR components

Finally, we set out to clarify underlying mechanisms attenuating the brain tumor targeting efficiency of heterogeneous ADCs. To investigate how each DAR component could affect biodistribution profiles, we prepared depatuxizumab-based MMAF ADCs with DARs of 4, 6, and 8 using our branched linkers and non-cleavable BCN–MMAF (see Supplementary Information for synthesis details). The parent N297A depatuxizumab was used as a DAR 0 control. These anti-EGFRvIII mAbs and conjugates were then labeled with Cy5.5 NHS ester at DOL of 2.3–2.5 to afford DAR 0 mAb **11**, DAR 4 ADC **12**, DAR 6 ADC **13**, and DAR 8 ADC **14** (**Figures 7A** and **S8**). Cy5.5 was installed directly onto the mAb scaffold so that the fluorescent signal would represent the localization of the entire ADC molecule. The fluorescent conjugates (3 mg/kg) were injected intravenously into NSG mice bearing orthotopic U87ΔEGFR-luc tumors on Day 5 post tumor implantation. After blood collection and cardiac perfusion at 48 h, major organs were harvested for fluorescence imaging. DAR 0 and DAR 4 conjugates **11** and **12** showed similar levels of brain tumor accumulation (**Figures 7B** and **7C**). In contrast, compared with DAR 4 ADC **12**, markedly attenuated brain tumor accumulation was observed for DAR 6 ADC **13** (*P* = 0.0396) and DAR 8 ADC **14** (*P* = 0.0288). Although ADCs **12**–**14** accumulated in the liver more significantly than DAR 0 mAb **11**, the degrees of liver accumulation and biodistribution patterns of these ADCs were similar and irrespective of DAR (**Figures 7D** and **7E**). In addition, the concentrations of DAR 4 and 6 ADCs **12** and **13** in blood were in a similar range and slightly below that of DAR 0 mAb **11** (**Figure 7F**). DAR 8 ADC **14** underwent accelerated clearance from the circulation probably because of greatly increased hydrophobicity. Collectively, these results demonstrate that high DAR components comprising a given heterogeneous ADC can show poor brain tumor targeting compared to components with optimal or low DARs, leading to reduced payload delivery efficiency.

**Figure 7.**
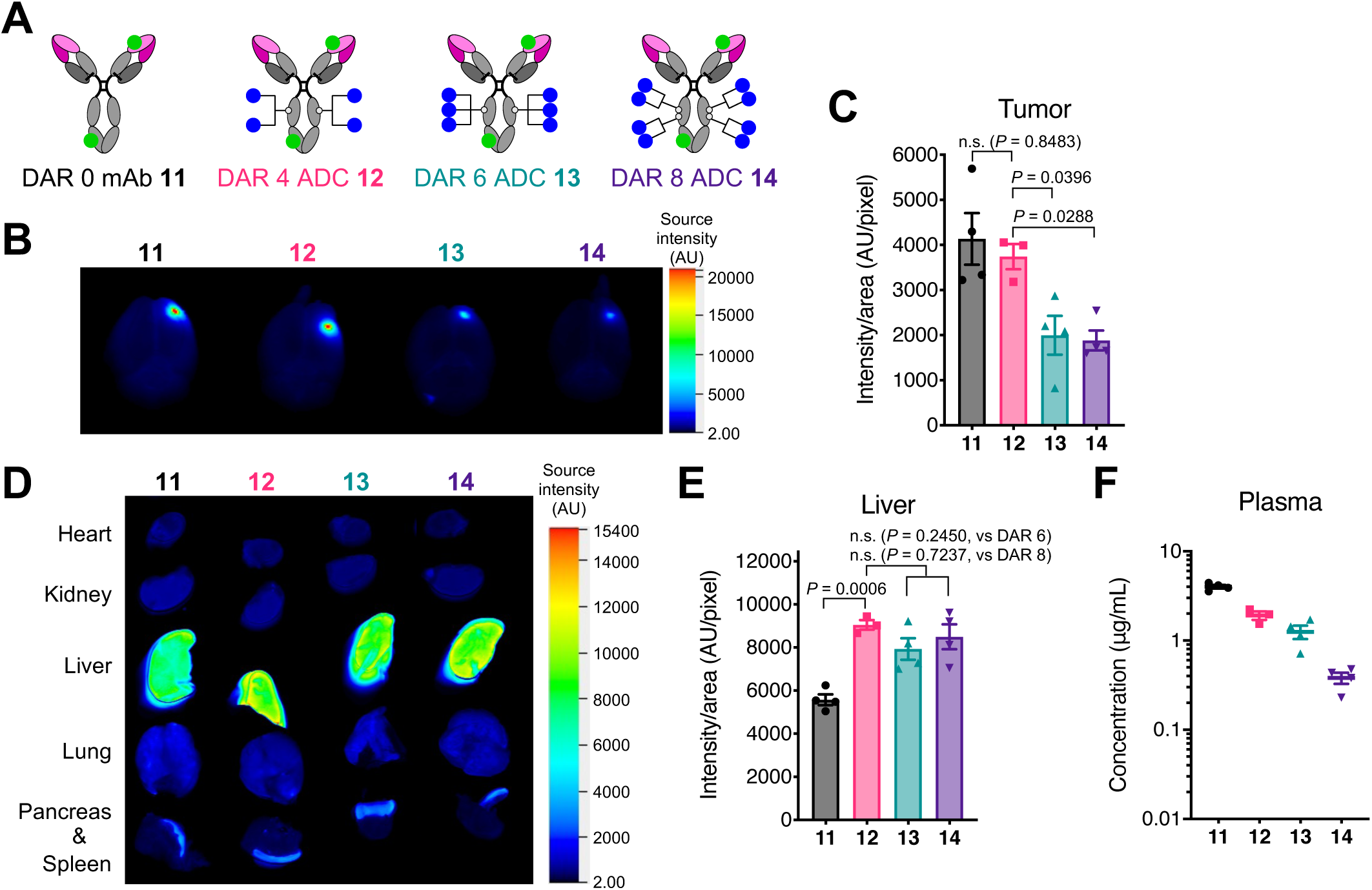
High-DAR components in heterogeneous ADCs target brain tumors less efficiently than components with optimal or low DAR. **A** Structures of fluorescently labeled anti-EGFRvIII ADCs equipped MMAF (blue circle) at DARs of 0, 4, 6, and 8. Cy5.5 (green circle) was conjugated by lysine coupling at DOL of 2.3–2.5. **B** Ex vivo fluorescence images of whole brains harvested from NSG mice bearing orthotopic U87ΔEGFR-luc tumors 48 hours after intravenous injection of each fluorescent ADC (n = 3 for DAR 4 ADC **12**; n = 4 for all other groups). Images were taken using a 700 nm channel. **C** Semi-quantification of the Cy5.5 signal derived from the tumor lesions in the whole brains. DAR 0 mAb **11** (black), DAR 4 ADC **12** (magenta), DAR 6 ADC **13** (green), and DAR 8 ADC **14** (purple). **D** Ex vivo fluorescence images of other major organs. **E** Semi-quantification of the Cy5.5 signal detected in the liver. **F** Concentrations in plasma. Blood was collected 48 h post ADC injection (right before cardiac perfusion) to quantify total antibody by sandwich ELISA. A representative image from each group is shown in all panels of fluorescence images. Data are presented as mean values ± SEM. For statistical analysis, a one-way ANOVA with a Dunnett’s post hoc test (control: DAR 4 ADC **12**) was used (see **Table S4** for details).

## DISCUSSION

We have investigated how ADC homogeneity impacts therapeutic efficacy and survival extension in orthotopic brain tumor models. Our stepwise conjugation method (i.e., MTGase-mediated site-specific linker conjugation and following click reaction) efficiently generated homogeneous ADCs targeting EGFR, EGFRvIII, and HER2. We tested these homogeneous ADCs and their heterogeneous variants prepared by conventional lysine or cysteine coupling for antiproliferative effect against brain tumor cells. In vitro, all DAR-matched ADCs showed comparable antigen-specific binding and cell killing potency irrespective of ADC homogeneity, conjugation method, or linker cleavability. However, we obtained contrasting results *in vivo*; all homogeneous ADCs exerted far better survival benefits in both cell line-derived xenograft and PDX orthotopic brain tumor models than could be achieved by corresponding heterogeneous variants, including a Depatux-M surrogate. Notably, a single dose of our homogeneous ADCs at 3 mg/kg provided complete remission in the orthotopic U87ΔEGFR-luc (2 out of 6 mice by anti-EGFR ADC **1**; 4 out of 6 mice by anti-EGFRvIII ADC **4**) and JIMT1-BR3 models (6 out of 6 mice by anti-HER2 ADC **6**), whereas DAR-matched heterogeneous ADCs did not in either case. To delve into this discrepancy, we performed biodistribution studies using intracranially xenografted GBM models. Our data from these studies indicate that homogeneous conjugation at optimal DARs likely allows for enhanced and persistent payload accumulation into intracranial tumors over several days, leading to improved *in vivo* efficacy. We also confirmed that both cleavable and non-cleavable linkers allowed homogeneous anti-EGFR conjugates to deliver payloads to intracranial GBM tumors at similar levels. Collectively, these results demonstrate that ADC homogeneity is a critical factor determining therapeutic efficacy in the treatment of brain tumors.

The question we asked next is how ADC homogeneity critically influences systemic payload delivery to brain tumors. Many studies have shown that homogeneous ADCs provide more favorable therapeutic effects in the treatment of other solid tumors than can be achieved by heterogeneous variants (Bryant et al., 2015; Junutula et al., 2008, 2010; Lhospice et al., 2015; Pillow et al., 2014). Nevertheless, the improvement in therapeutic efficacy observed in our study appears to be much more prominent compared to those cases. We think that blockage of drug influx by an intact BBB in and around brain tumors likely answers this question. The BBB was believed to be uniformly and significantly disrupted in most GBM tumors. Contrary to this previous belief, recent preclinical and clinical studies have demonstrated that a measurable number of GBM cells, in particular ones near the growing edge of the infiltrative tumor area, exist behind an intact BBB or partially functional blood– tumor barrier (BTB) (Arvanitis et al., 2020; Kim et al., 2018; Marin et al., 2021; Sarkaria et al., 2018; van Tellingen et al., 2015). As such, GBM cells protected by an intact BBB are inaccessible to systemically administered ADCs. Recently, Sarkaria and co-workers exhaustively validated heterogeneous BBB disruption in multiple PDX models of GBM, including GBM12 (Marin et al., 2021). They also demonstrated that the intact BBB likely caused an uneven intracranial distribution of systemically administered Depatux-M, resulting in insignificant treatment outcomes in 5 out of 7 orthotopic PDX models. In contrast, they found that Depatux-M exerted remarkable therapeutic effects when tested in subcutaneous models of the same PDX tumors, in which the BBB did not constitute the tumor microenvironment. This report highlights the importance of testing ADCs for brain tumor treatment in clinically relevant orthotopic models rather than subcutaneous models.

Our findings and the report from the Sarkaria group (Marin et al., 2021) lead to a hypothesis that high-DAR species included in heterogeneous ADCs cannot be efficiently delivered to intracranial tumors across the BBB compared to low-DAR species. Consequently, the effective DAR (i.e., DARs adjusted based on the brain tumor targeting efficiency of each DAR component relative to that of the unmodified mAb) and payload dose are considerably reduced (**Figure 8**). In contrast, homogeneous ADCs constructed at optimal DARs likely undergo only marginal impairment in brain tumor targeting, leading to a minimal reduction in effective payload dose. Indeed, our intravital imaging study showed that the difference in payload dose between heterogeneous and homogeneous ADCs could reach up to 2.5-fold. In general, ADC hydrophobicity increases in proportion to the degree of payload conjugation. As such, high-DAR ADCs have greater aggregation tendency compared with low-DAR ADCs. In circulation, such multimolecular complexation may also occur with abundant proteins such as albumin, resulting in increased apparent hydrodynamic radius (Frka-Petesic et al., 2016). Considering that BBB permeability declines exponentially with molecular size (Li et al., 2016), we speculate that an increase in apparent hydrodynamic radius is disadvantageous for delivering conjugated payloads to brain tumors across the intact BBB or partially functional BTB. As demonstrated in our treatment study using the intracranial JIMT-1-BR3 tumor model (i.e., complete remission in all animals by homogeneous ADC **6** vs no complete remission by heterogeneous ADC **7**), a decrease in the effective DAR by heterogeneous conjugation could be further prominent in grade 1–3 gliomas and HER2-positive brain metastatic tumors, in which BBB disruption is less significant than in GBM (Gril et al., 2020; Yonemori et al., 2010). The use of more hydrophobic payloads than MMAF may also make this effect salient. As observed in previous studies using other solid tumor models (Hamblett et al., 2004; Lhospice et al., 2015), clearance and *in vivo* stability of ADCs could also be factors influencing payload delivery efficiency and overall treatment efficacy in orthotopic brain tumor models. Indeed, we observed promoted clearance for DAR 8 MMAF ADC **14**. However, DAR 6 ADC **13**, which also showed poor brain tumor targeting, did not undergo rapid clearance. In addition, the treatment efficacy of heterogeneous Cys conjugate **2** in the orthotopic U87ΔEGFR-luc tumor model was comparable to that of Lys conjugate **3**, despite its impaired thermal and circulation stability. Overall, these findings support the conclusion that ADC homogeneity can influence payload delivery to brain tumors across the BBB more significantly than clearance and *in vivo* stability profiles. Future in-depth structural and mechanistic studies will clarify the validity of our hypothesis in other combinations of mAbs, linker and conjugation chemistries, and payload types.

**Figure 8.**
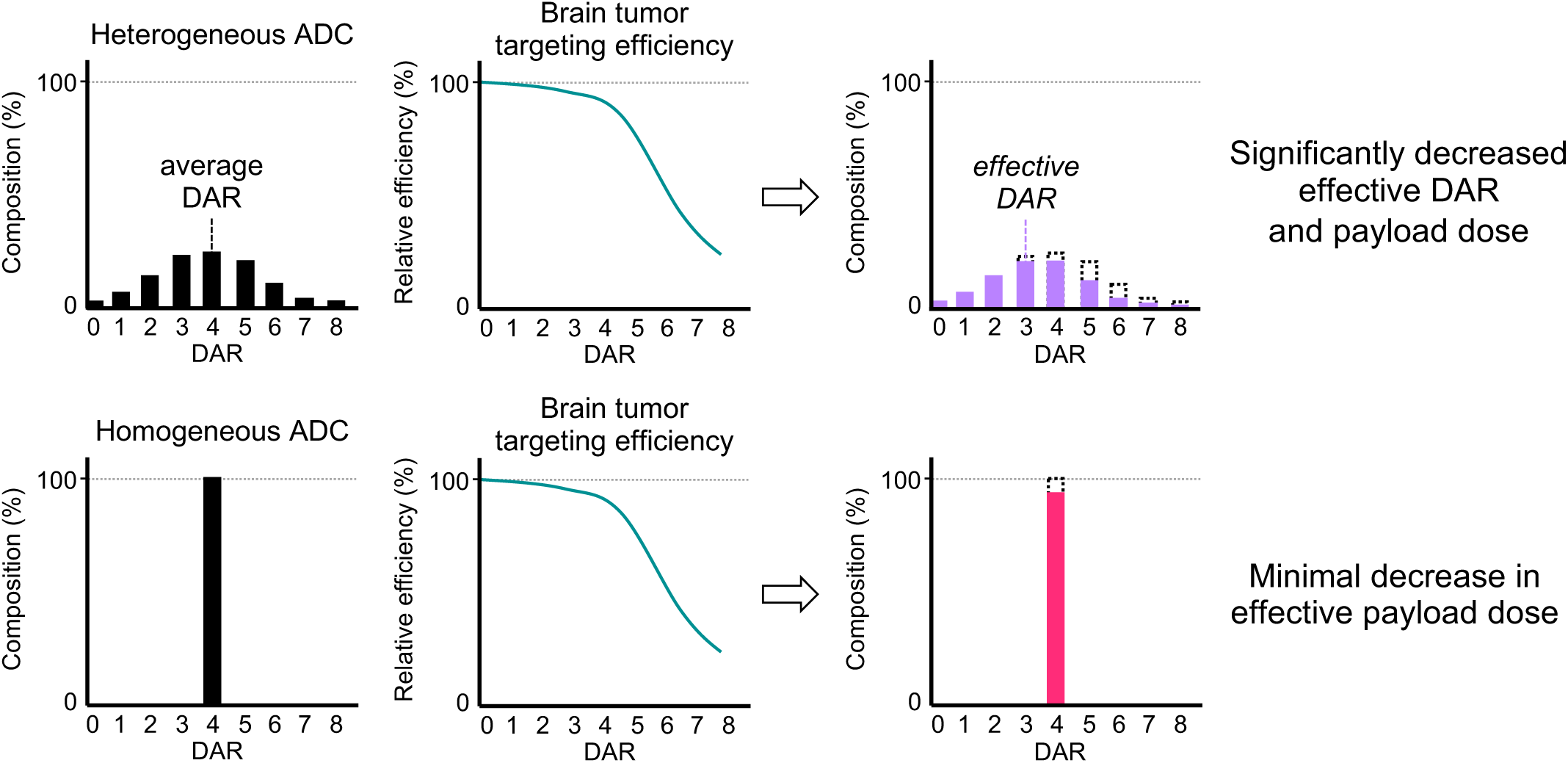
Reduction in effective DAR and payload dose is more prominent in heterogeneous ADCs than in homogeneous ADCs. All values used in this figure are estimated values based on the data shown in **Figures 1** and **7**. Theoretical payload doses of heterogeneous and homogeneous ADCs with the same (average) DAR are equivalent if administered at the same mAb dose. However, high-DAR components in heterogeneous ADCs show poor brain tumor targeting, decreasing the effective DAR and payload dose. Such deterioration is marginal in the case of homogeneous ADCs, leading to improved payload delivery and overall efficacy.

In summary, our findings highlight the critical importance of ADC homogeneity in maximizing efficacy in brain tumor treatment. Employing homogeneous conjugation at optimal DARs with properly designed linkers could be a promising approach to resurrecting the ADCs for GBM that have failed to show therapeutic benefits in clinical trials, including Depatux-M. We also envision that initiating exploration of new ADCs using homogeneous conjugation technologies will help streamline the optimization of ADC properties (e.g., DAR, hydrophobicity, stability) and effectively expand our repertoire of promising drug candidates for brain tumors. In addition to this updated molecular design guideline, further understanding of brain tumor biology and pathophysiology will also be crucial to identify promising combinations of antibody targets, ADC linker properties (e.g., structure, drug release mechanism), and payload types. In particular, deeper understanding of the integrity and functions of the BBB found in patient-derived brain tumor samples could open up the next step to improving payload delivery efficiency. We believe that such multifaceted approaches will finally lead us to novel ADCs or other targeted therapy modalities with the potential to conquer GBM and other intractable brain tumors.

## ACKNOWLEDGEMENTS

We gratefully acknowledge the following researchers for providing the cells used in this study: U87ΔEGFR from Dr. Erwin G. Van Meir (Emory University), Gli36δEGFR from Dr. E. Antonio Chiocca (Brigham and Women’s Hospital), JIMT-1-BR3 from Dr. Patricia S. Steeg (the National Cancer Institute), and GBM12 from Dr. Jann N. Sarkaria (Mayo Clinic). We thank Dr. Chisato M. Yamazaki for providing scientific input and Mr. Travis J. Roeder for technical assistance. This work was supported by the National Institutes of Health (R35GM138264 to K.T.; R61NS112410 and P01CA163205 to B.K.), the Department of Defense Breast Cancer Research Program (W81XWH-18-1-0004 and W81XWH-19-1-0598 to K.T.), the Cancer Prevention and Research Institute of Texas (RP150551 and RP190561 to Z.A.), the Welch Foundation (AU-0042-20030616 to Z.A.), and the Japan Society for the Promotion of Science (postdoctoral fellowship to Y.A. and A.Y.).

## AUTHOR CONTRIBUTIONS

Conceptualization, K.T.; Methodology, Y.A., Y.O., W.X., N.Z., Z.A., B.K., and K.T.; Validation, Y.A., Y.O., and W.X.; Formal Analysis, Y.A., Y.O., S.Y.Y.H; Investigation, Y.A., Y.O., W.X., S.Y.Y.H, A.Y., and K.T.; Resources, N.Z., Z.A., B.K., and K.T.; Writing – Original Draft, Y.A., Y.O., and K.T.; Writing – Review & Editing, Y.A., Y.O., A.Y., Z.A., and K.T.; Visualization, Y.A. and Y.O.; Supervision, K.T.; Project Administration, K.T.; Funding Acquisition, Y.A., A.Y., Z.A., B.K., and K.T.

## METHODS

### Compounds and antibody conjugates

See Supplementary Information for synthesis details and characterization data of all compounds used in this study.

### Antibodies

Anti-EGFR, anti-EGFRvIII, and anti-HER2 IgG1 mAbs with N88A/N297A, N297A, or N297Q mutation were expressed in-house (see below). The other antibodies used in this study were purchased from commercial vendors as follows: Rabbit anti-MMAF antibody (LEV-PAF1) from Levena Biopharma; goat anti-human IgG Fab-horseradish peroxidase (HRP) conjugate (109-035-097), goat anti-human IgG Fc antibody (109-005-098), and donkey anti-human IgG-HRP conjugate (709-035-149) from Jackson ImmunoResearch; goat anti-rabbit IgG–HRP conjugate (32260) from Thermo Fisher Scientific; rabbit anti-cleaved caspase 3 antibody (9661S) and rabbit anti-EGFR antibody (4267S) from Cell Signaling Technology); and rabbit anti-Ki67 antibody (ab16667) from Abcam.

### Expression and purification of human monoclonal antibodies

All human monoclonal antibodies were produced according to the procedure reported previously(Anami et al., 2017; Shi et al., 2014). Briefly, free style HEK-293 human embryonic kidney cells (Invitrogen) were transfected with a mammalian expression vector encoding for the human IgG1 kappa light chain and full-length heavy chain sequences (based on variable sequences of cetuximab, depatuxizumab, or trastuzumab). The transfected HEK-293 cells were cultured in a humidified cell culture incubator at 37 °C with 8% CO_2_ and shaking at 150 rpm for 7 days before harvesting the culture medium. The antibody secreted into the culture medium was purified using Protein A resin (GE Healthcare).

### MTGase-mediated antibody–linker conjugation

Anti-EGFR mAb with N88A/N297A double mutations (400 µL in PBS, 5.53 mg/mL, 2.21 mg antibody) was incubated with the diazide branched linker developed by us previously (Anami et al., 2017, 2018) (5.9 µL of 100 mM stock in water, 40 equiv.) and Activa TI^®^ (101 µL of 40% solution in PBS, Ajinomoto, purchased from Modernist Pantry) at room temperature for 22 h. The reaction was monitored using either 1) an Agilent LC-MS system consisting of a 1100 HPLC and a 1946D single quadrupole ESI mass spectrometer equipped with a MabPac RP column (3 × 50 mm, 4 µm, Thermo Scientific) or 2) a Thermo LC-MS system consisting of a Vanquish UHPLC and a Q Exactive™ Hybrid Quadrupole-Orbitrap(tm) Mass Spectrometer equipped with a MabPac RP column (2.1 × 50 mm, 4 µm, Thermo Scientific). Elution conditions were as follows: mobile phase A = water (0.1% formic acid); mobile phase B = acetonitrile (0.1% formic acid); gradient over 6.8 min from A:B = 75:25 to 1:99; flow rate = 0.5 mL/min. The conjugated antibody was purified by SEC (Superdex 200 increase 10/300 GL, GE Healthcare, solvent: PBS, flow rate = 0.6 mL/min) to afford an antibody–linker conjugate [1.91 mg, 86% yield determined by bicinchoninic acid (BCA) assay].

### Construction of homogeneous ADCs by strain-promoted azide-alkyne cycloaddition

BCN–EVCit–PABC–MMAF (20.7 µL of 3.7 mM stock solution in DMSO, 1.5 equivalent per azide group) was added to a solution of the mAb–linker conjugate in PBS (460 µL, 4.16 mg/mL), and the mixture was incubated at room temperature for 22 h. The reaction was monitored using either Agilent LC-MS system or Thermo LC-MS system equipped with a MabPac RP column (see above) and the crude products were purified by SEC to afford homogeneous ADC **1** (1.71 mg, 90% yield determined by BCA assay). Analysis and purification conditions were the same as described above. Homogeneity was confirmed by ESI-MS analysis. Homogeneous anti-EGFRvIII ADC **4** and anti-HER2 ADC **6** were prepared in the same manner.

### Construction of a heterogeneous ADC by cysteine conjugation

Aglycosylated anti-EGFR mAb (298 µL in PBS, 3.0 mg/mL, 895 µg antibody) was mixed with TCEP (19.1 µL of 1 mM stock solution in water, 3.2 equiv.) and EDTA (30 µL of 10 mM stock solution in water, pH 9, 10% v/v) and incubated at 37 °C for 2 h. MC–MMAF (9.0 µL of 10 mM stock solution in DMSO, 15 equiv.) was added to the partially reduced mAb solution and the reaction mixture was incubated overnight at room temperature. The reaction was monitored using an Agilent 1100 HPLC system equipped with a MAbPac HIC-Butyl column (4.6 × 100 mm, 5 µm, Thermo Scientific). Elution conditions were as follows: mobile phase A = 50 mM sodium phosphate containing ammonium sulfate (1.5 M) and 5% isopropanol (pH 7.4); mobile phase B = 50 mM sodium phosphate containing 20% isopropanol (pH 7.4); gradient over 25 min from A:B = 99:1 to 1:99; flow rate = 0.8 mL/min. *N*-acetyl cysteine (4.5 µL of 100 mM stock solution in DMSO, 75 equiv.) was added to the reaction mixture for quenching the reaction. The crude products were purified by SEC to afford Cys conjugate **2** (668 µg, 75% yield determined by BCA assay, average DAR: 3.8). SEC purification conditions were the same as described above. The average DAR value was determined based on UV peak areas in HIC analysis.

### Construction of heterogeneous ADCs by lysine conjugation

Aglycosylated anti-EGFR mAb (105 µL in PBS, 3.0 mg/mL, 315 µg antibody) was mixed with 1 M phosphate solution at pH 9 (10.5 µL) and MMAF-NHS (2.5 µL of 10 mM stock solution in DMSO, 12 equiv.) and the mixture was incubated at room temperature for 3 h. The reaction was monitored using either Agilent LC-MS system or Thermo LC-MS system equipped with a MabPac RP column (see above). The crude products were purified by SEC to afford Lys conjugate **3** (197 µg, 63% yield determined by BCA assay, average DAR: 3.9). Analysis and purification conditions were the same as described above. The average DAR value was determined based on ion intensity of each DAR species in ESI-MS analysis. Heterogeneous anti-EGFRvIII ADC **5** and anti-HER2 ADC **7** were constructed in the same manner.

### Construction of anti-EGFR Cy5.5 conjugates

Cy5.5 conjugates **8–10** were prepared in the same manner as the preparation of corresponding ADCs described above. Instead of MMAF-containing linker modules, either of the following linker modules were used: DBCO–EVCit–Cy5.5 (synthesized in house, for homogeneous Cy5.5 conjugate **8**), Cy5.5 maleimide (purchased from Click Chemistry Tools, for Cys-Cy5.5 conjugate **9**), Cy5.5-NHS ester (purchased from Click Chemistry Tools, for Lys-cy5.5 conjugate **10**), or DBCO–Cy5.5 (purchased from Click Chemistry Tools, for homogeneous non-cleavable Cy5.5 conjugate). Degrees of labeling (DOL) were determined by ESI-MS analysis (based on ion intensity of each DOL species) or using a plate reader (BioTek Synergy HTX) with a standard curve for free Cy5.5 (absorbance at 680 nm).

### Construction of anti-EGFRvIII MMAF-Cy5,5 conjugates

Homogeneous anti-EGFRvIII MMAF ADCs with DARs of 4, 6, and 8 were prepared from depatuxizumab with an N297A (for DAR 4 and 6) or N297Q mutation (for DAR 8). For the preparation of the DAR 6 MMAF ADC, the diazido-methyltetrazine tri-arm linker developed by us previously (Yamazaki et al., 2021) was used. Subsequently, unmodified N297A depatuxizumab (DAR 0) and each ADC were labeled with Cy5.5-NHS ester (10 mM stock solution in DMSO, 6–8 equiv.) to achieve an average DOL of 2.3–2.5. The labeling reaction was performed in the same manner as described above, except that the reaction was quenched with ethanol amine (100 mM stock solution in water, 20 equiv.). The average DOL values of MMAF-Cy5.5 conjugates **11**–**14** were determined based on ion intensity of each DOL species in ESI-MS analysis.

### Long-term stability test

Each ADC (1 mg/mL, 10 µL) in PBS was incubated at 37 °C for 28 days and stored at –80 °C until use. Samples were analyzed using an Agilent 1100 HPLC system equipped with a MAbPac SEC-1 analytical column (4.0 × 300 mm, 5 µm, Thermo Scientific). The conditions were as follows: flow rate = 0.2 mL/min; solvent = PBS. All assays were performed in triplicate.

### Cell lines

U87ΔEGFR was received from Dr. Erwin G. Van Meir (Emory University). Gli36δEGFR was received from Dr. E. Antonio Chiocca (Brigham and Women’s Hospital, Harvard Medical School). U87ΔEGFR-luc was generated by lentiviral transduction of U87ΔEGFR cells using Lentifect™ lentiviral particles encoding for firefly luciferase and a puromycin-resistant gene (GeneCopoeia, LP461-025). Transduction was performed according to the manufacturer’s instruction. U87ΔEGFR, U87ΔEGFR-luc, Gli36δEGFR, and HEK293 (ATCC) cells were cultured in DMEM (Corning) supplemented with 10% EquaFETAL^®^ (Atlas Biologicals), GlutaMAX^®^ (2 mM, Gibco), and penicillin-streptomycin (penicillin: 100 units/mL; streptomycin: 100 µg/mL, Gibco). JIMT1-BR3 was received from Dr. Patricia S. Steeg (National Cancer Institute) and maintained in RPMI1640 (Corning) supplemented with 10% EquaFETAL^®^, GlutaMAX^®^ (2 mM), sodium pyruvate (1 mM, Corning), and penicillin-streptomycin (penicillin: 100 units/mL; streptomycin: 100 µg/mL). GBM12 was received from Dr. Jann N. Sarkaria (Mayo Clinic). RFP-expressing GBM12 (GBM12-RFP) was generated by transduction with lentivirus (System Biosciences, LL110VA-1) according to the manufacturer’s instruction. GBM12 and GBM12-RFP cells were maintained in DMEM supplemented with 2% fetal bovine serum and penicillin-streptomycin (penicillin: 100 units/mL; streptomycin: 100 µg/mL). All cells except U87ΔEGFR-luc and HEK293 were authenticated via short tandem repeat profiling before use. All cells were cultured at 37 °C under 5% CO_2_, and passaged before becoming fully confluent up to 40 passages. All cells were periodically tested for mycoplasma contamination.

### Cell-based ELISA

Cells (U87ΔEGFR or HEK293) were seeded in a culture-treated 96-well clear plate (10,000 cells/well in 100 μL culture medium) and incubated at 37 °C with 5% CO_2_ for 24 h. Paraformaldehyde (8%, 100 μL) was added to each well and incubated for 15 min at room temperature. The medium was discarded and the cells were washed three times with 100 μL of PBS. Cells were treated with 100 μL of blocking buffer (0.2% BSA in PBS) with agitation at room temperature for 2 h. After the blocking buffer was discarded, serially diluted samples (in 100 µL PBS containing 0.1% BSA) were added and the plate was incubated overnight at 4 °C with agitation. The buffer was discarded and the cells were washed three times with 100 μL of PBS containing 0.25% Tween 20. Cells were then incubated with 100 μL of donkey anti-human IgG–HRP conjugate (diluted 1:10,000 in PBS containing 0.1% BSA) was added and the plate was incubated at room temperature for 1 h. The plate was washed three times with PBS containing 0.25% Tween 20, and 100 μL of 3,3’,5,5’-tetramethylbenzidine (TMB) substrate (0.1 mg/mL) in phosphate–citrate buffer/30% H_2_O_2_ (1:0.0003 volume to volume, pH 5) was added. After color was developed for 10-30 min, 25 μL of 3 N-HCl was added to each well and then the absorbance at 450 nm was recorded using a plate reader (BioTek Synergy HTX). Concentrations were calculated based on a standard curve. K_D_ values were then calculated using Graph Pad Prism 8 software. All assays were performed in triplicate.

### Cell viability assay

Cells were seeded in a culture-treated 96-well clear plate (5,000 cells/well in 50 μL culture medium) and incubated at 37 °C under 5% CO_2_ for 24 h. Serially diluted samples (50 µL) were added to each well and the plate was incubated at 37 °C for 72 h. After the old medium was replaced with 100 µL fresh medium, 20 μL of a mixture of WST-8 (1.5 mg/mL, Cayman chemical) and 1-methoxy-5-methylphenazinium methylsulfate (100 μM, Cayman Chemical) was added to each well, and the plate was incubated at 37 °C for 2 h. After gently agitating the plate, the absorbance at 460 nm was recorded using a plate reader (BioTek Synergy HTX). EC_50_ values were calculated using Graph Pad Prism 8 software. All assays were performed in triplicate.

### Animal studies

All procedures were approved by the Animal Welfare Committee of the University of Texas Health Science Center at Houston and performed in accordance with the institutional guidelines for animal care and use. All animals were housed under controlled conditions, namely 21–22 °C (+/-0.5 °C), 30– 75% (+/-10%) relative humidity, and 12:12 light/dark cycle with lights on at 7.00 a.m. Food and water were available ad libitum for all animals. NSG mice were purchased from The Jackson Laboratory (stock number: 005557) and bred in house. CD-1^®^ mice was purchased from Charles River Laboratories (Strain Code: 022) and used without in-house breeding.

### Orthotopic xenograft mouse models of human brain tumors

U87ΔEGFR-luc (1 × 10^5^ cells), GBM12 (2 × 10^5^ cells), or JIMT1-BR3 (2 × 10^5^ cells) were stereotactically implanted into NSG mice (6–8 weeks old, male and female) based on the previously reported method (Otani et al., 2020). Typical procedure. NSG mice were injected intraperitoneally with a cocktail of ketamine (67.5 mg/kg) and dexmedetomidine (0.45 mg/kg) and maintained at 37 °C on a heating pad until the completion of surgery. After the head skin was shaved and treated with 10 μL of 0.25% bupivacaine supplemented with epinephrine (1:200,000), anesthetized mice were placed on a stereotactic instrument. After disinfecting the head skin with chlorhexidine and ethanol, a small incision was made and then a burr hose was drilled into the skull over the right hemisphere (1 mm anterior and 2 mm lateral to the bregma). A 10 μL Hamilton syringe (model 701 N) was loaded with cells suspended in 2 μL cold hanks-balanced salt solution (HBSS) and slowly inserted into the right hemisphere through the burr hole (3.5 mm depth). After a 1-min hold time, cells were injected over a 5-min period (0.4 μL/min). After a 3-min hold time, the needle was retracted at a rate of 0.75 mm/min. The incision was closed using GLUture^®^ (Zoetis) and mice were injected with atipamezole (1 mg/kg, i.p.).

### Treatment study

Brain tumor-bearing NSG mice were randomized and injected intravenously with a single dose of either ADC (3 mg/kg) or PBS. Group assignment and dose schedule were as follows: U87ΔEGFR-luc model, n = 4 or 6 for vehicle, n = 6 for ADCs, injected on Day 5; GBM12 model, n = 15 for vehicle, n = 14 for ADCs injected on Day 8; JIMT-1-BR3 model, n = 6 for all groups, injected on Day 7. Growth of U87ΔEGFR-luc tumors was monitored by bioluminescence imaging (BLI) using an Xtreme *in vivo* imager (Bruker Biospin, upper limit: 1.5 × 10^5^ photons/sec/mm^2^; lower limit: 5.0 × 10^3^ photons/sec/mm^2^) once every week. Tumor growth was also evaluated by MRI (see the following sections for details). Body weight was monitored every 3–4 days and mice were euthanized when body weight loss of >20% or any severe clinical symptom was observed.

### MRI and measurement of tumor volume (GBM12 model)

MRI was performed using a 7 Tesla MRI scanner (Bruker Biospin) on Day 18 post tumor implantation. Tumor-bearing mice (n = 4/group, randomly selected from each group) were anesthetized with 1.5% isoflurane in a 30:70 mixture of O_2_ and medical air. MRI contrast agent (Dotarem) was injected (50 µL, i.p.) before imaging to help visualize the tumor. T2-weighted images were acquired using a multi-echo RARE sequence with a RARE factor of 3. Acquisition parameters were as follows: TR = 5000 ms, TE = 17 42.5 68 and 93.5 ms, 15 image slices with 100 µm slice thickness, in-plane resolution = 100 × 100 µm^2^. ImageJ software was utilized to measure the tumor volume. Regions of interest (ROI) were manually drawn to circumscribe the entire tumor, and volume was calculated by counting all the voxels within the ROI and multiplying the total number of pixels by the volume of the voxel (100 × 100 × 500 µm^3^).

### MRI in the U87ΔEGFR-luc and JIMT-1-BR3 models

MRI images were taken using a 7 Tesla MRI scanner (Bruker Biospin) on Day 58 (JIMT-1-BR3) or Day 61 (U87ΔEGFR-luc) post tumor implantation. Tumor-bearing mice (U87ΔEGFR-luc model: 4 survivor mice treated with homogeneous anti-EGFRvIII ADC **4**; JIMT-1: n = 6/group) were anesthetized with 2% isoflurane throughout the imaging procedure. A 35mm ID volume coil (Bruker Biospin) receive setup was used for data acquisition. T2-weighted coronal and axial images were acquired with a Spin Echo RARE sequence. Acquisition parameters were as follows: TR = 3000 ms, TE =57 ms, RARE factor 12, 6 NAV, Slice thickness of 0.75 mm, slice gap 0.25 mm, in plane resolution of 156 μm for coronal and 117 μm for axial.

### Immunohistochemistry

Mice were euthanized at the end of the treatment study in the GBM12 model and their excised tumor-bearing brain were embedded in paraffin. Samples were deparaffinized using xylene and rehydrated in decreasing concentration of ethanol. Subsequently, slices were incubated in 0.3% H_2_O_2_ for 30 min and autoclaved for 15 min at 121°C in citrate buffer. After blocking with animal-free blocking solution, slices were incubated with either rabbit anti-cCaspase 3 antibody (1:250), rabbit anti-EGFR antibody (1:50), or rabbit anti-ki67 antibody (1:200). SignalStain^®^ Boost IHC Detection Reagent and DAB substrate kit (Cell Signaling Technology) were used and then the sections were counterstained with hematoxylin. Bright-field images were taken using an EVOS-FL Auto2 imaging system (Invitrogen). For cleaved caspase-3 and ki67 quantification, three representative areas of each stained sample were imaged and the populations of cCaspase3- and ki67-positive cells were analyzed using Image J software.

### In vivo pharmacokinetic study

CD-1^®^ mice (6–8 weeks old, female, n = 3/group) were injected intravenously with each mAb or ADC (3 mg/kg). Blood samples (5 µL) were collected from each mouse via the tail vein at each time point (15 min, 6 h, 1 day, 2 days, 4 days, 9 days, and 14 days) and immediately processed with 495 µL of 5 mM EDTA/PBS. After removal of cells by centrifugation (10 min at 10,000 × *g* at 4 °C), plasma samples were stored at –80 °C until use. All mice were humanely killed after last blood collection. Plasma samples were analyzed by sandwich ELISA. For determination of the total antibody concentration (both conjugated and unconjugated), a high-binding 96-well plate (Corning) was treated with goat anti-human IgG Fc antibody (500 ng/well). After overnight surface coating at 4 °C, the plate was blocked with 100 µL of 2% BSA in PBS containing 0.05% Tween 20 (PBS-T) with agitation at room temperature for 1 h. Subsequently, the solution was removed and each diluted plasma sample (100 µL, diluted with PBS-T containing 1% BSA) was added to each well, and the plate was incubated at room temperature for 2 h. After each well was washed three times with 100 µL of PBS-T, 100 µL of goat anti-human IgG Fab– HRP conjugate (1:5,000) was added. After being incubated at room temperature for 1 h, the plate was washed and color development was performed as described above (see the section of “Cell-based ELISA”) . For determination of ADC concentration (conjugated only), assays were performed in the same manner using the following proteins and antibodies: human EGFR (100 ng/well, #EGR-H5222 from ACROBiosystems) for plate coating, and rabbit anti-MMAF antibody (1:5,000) and goat anti-rabbit IgG–HRP conjugate (1:10,000) as secondary and tertiary detection antibodies, respectively.

Concentrations were calculated based on a standard curve. Half-life at the elimination phase (t_1/2β_, day) and clearance rate [CL, (mg/kg)/(μg/mL)/day] of each conjugate were estimated using methods for non-compartmental analysis (Gabrielsson and Weiner, 2012). PKSolver (a freely available menu-driven add-in program for Microsoft Excel) (Zhang et al., 2010) was used for this calculation. Area under the curve (AUC_0-14 days_, μg/mL × day) was calculated using GraphPad Prism 8 software. See **Table S3** for all observed PK parameters.

### Ex vivo fluorescence imaging and quantification

Intracranial U87ΔEGFR-luc tumor-bearing NSG mice (6-8 weeks old, male and female) were prepared as described above and randomized into three groups (n = 3) 5 days post tumor implantation. Each Cy5.5 conjugate was administered intravenously at 3 mg/kg. After 48 h, the tumor-bearing mice were anesthetized with ketamine/xylazine. Subsequently, the mice underwent cardiac perfusion with PBS(+) containing sodium heparin (10 units/mL) and 4% paraformaldehyde/PBS(+). Major organs including the brain were then harvested. Cy5.5-based near-infrared fluorescence images of the harvested organs were taken using a LI-COR Odyssey 9120 imager (Ex: 685 nm laser, intensity: L1.0 for brain, L2.0 for other organs, Em: 700 nm channel). Semi-quantification of the signals from ROIs was also performed using LI-COR Image Studio software. For tissue imaging, the brain samples were embedded in paraffin and tissue sections were prepared (thickness: 5 µm). After de-paraffinization of using toluene, mounting medium containing DAPI (VECTOR #H-1200) was applied to the tissue slides. Fluorescence Images were taken using a Nikon Eclipse TE2000E inverted microscope (Cy5 channel). Three ROIs in each sample were acquired and analyzed for semi-quantification using ImageJ software.

### Intravital imaging

Male NSG mice (6–8 weeks old) were implanted with GBM12-RFP cells (2 × 10^5^ cells) stereotactically into the right hemisphere (2 mm lateral and 2.5 mm posterior to bregma, 1 mm depth) as previously described (Nair et al., 2020). Thirteen days after tumor implantation, craniectomy was performed over the tumor-implanted area. Cover glass (Bioscience Tools) was placed on the brain surface and glued to the skull with dental resin. Next day (Day 14), each Cy5.5 conjugate was administrated intravenously at 3 mg/kg (n = 4 for Cys-Cy5.5 conjugate **9**; n = 3 for the other groups). For intravital imaging, mice were anesthetized with isoflurane and positioned on the stage of a A1R-MP confocal microscope (NIKON) equipped with ×16 water immersion objective lens. Subsequently, 100 μL of 2% FITC-conjugated dextran (500 kDa, Sigma) was administrated through the tail vein, and Z-stack images were acquired based on Cy5.5, FITC, and RFP signals. Pre- and post-treatment images were acquired on Day 14, 15, 17, 19, and 22 after tumor implantation. The images were analyzed using NIS Elements AR software (NIKON). Intensity of the RFP (derived from GBM12-RFP) and Cy5.5 signals in two or three independent ROIs were calculated to determine Cy5.5/GBM12-RFP ratios.

### Statistical Analyses

Kaplan-Meier survival curve statistics were analyzed with a log-rank (Mantel–Cox) test. To control the family-wise error rate in multiple comparisons, crude *P* values were adjusted by the Bonferroni method. Differences with adjusted *P* values <0.05 were considered statistically significant in all analysis. For immunohistochemistry, immunofluorescence, ex vivo fluorescence imaging, MRI, and intravital imaging, a one-way ANOVA with a Tukey–Kramer or Dunnett’s post hoc test was used for multiple comparisons. See **Table S4** for all *P* values.

