## Supplemental File for "Homogeneity of antibody-drug conjugates critically impacts the therapeutic efficacy in brain tumors"

### Supplementary Figures

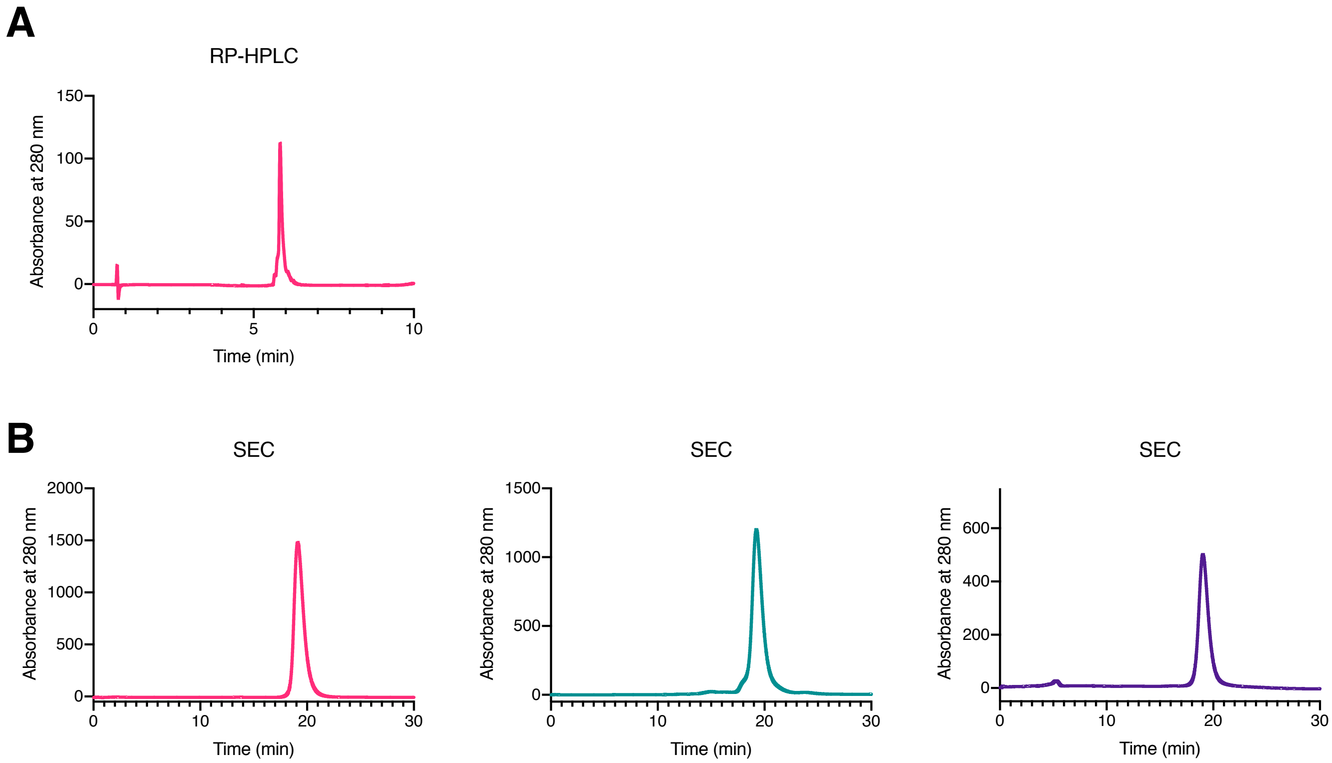


**Figure S1.** Reverse-phase HPLC and size-exclusion chromatography (SEC). (A) Reverse-phase HPLC trace of homogeneous ADC **1** before SEC purification (UV absorbance: 280 nm). The average DAR was determined to be 4 based on the lack of lower DAR species. (B) SEC traces (UV absorbance: 280 nm) of homogeneous ADC **1** (magenta), Cys-conjugate **2** (green), and Lys-conjugate **3** (purple).

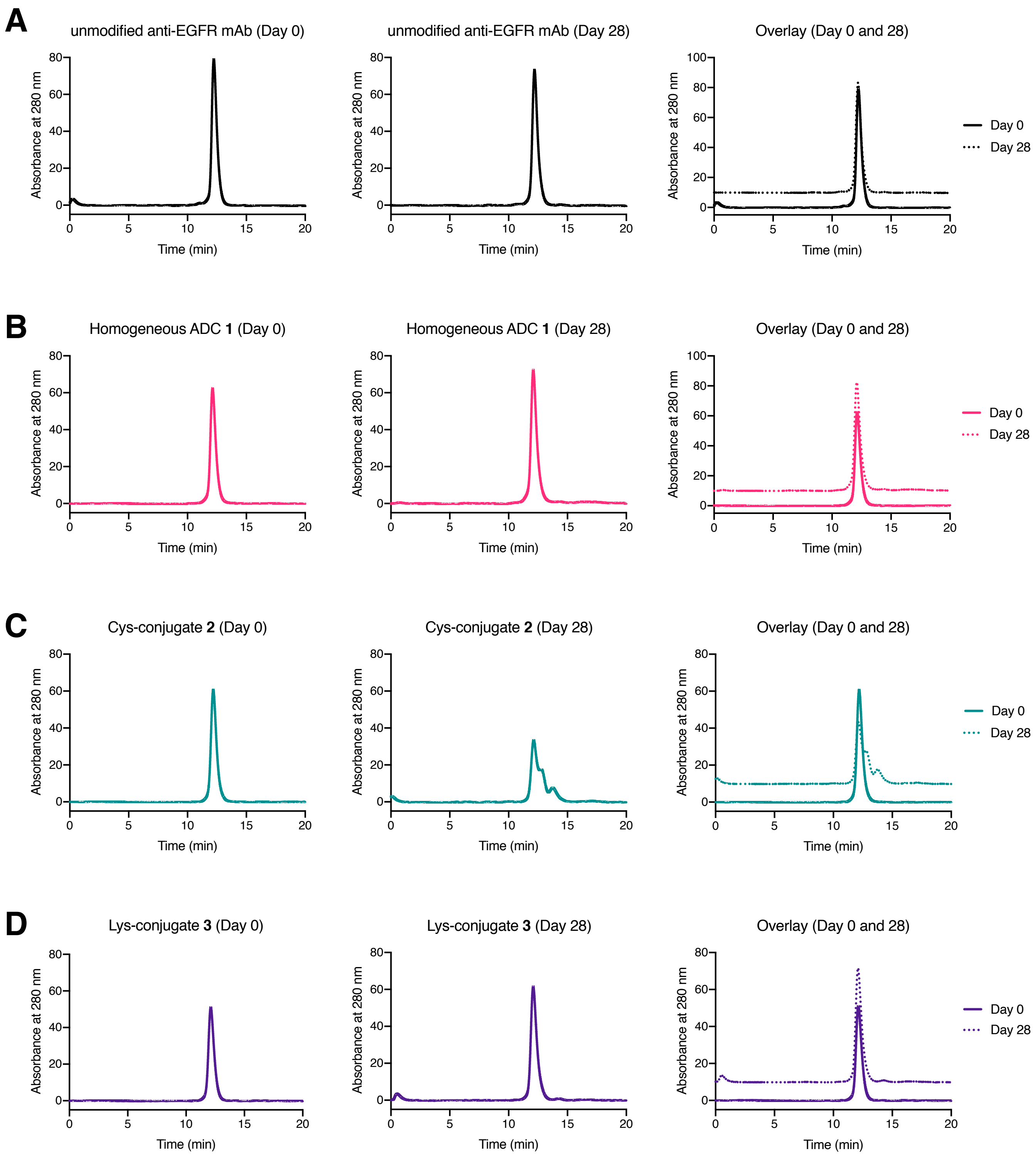


**Figure S2.** SEC analysis of ADCs after 1-month incubation at 37 ºC in PBS (pH 7.4). (A) Aglycosylated anti-EGFR mAb (cetuximab mutant), (B) homogeneous ADC **1**, (C) Cys-conjugate **2**, and (D) Lys-conjugate **3**.

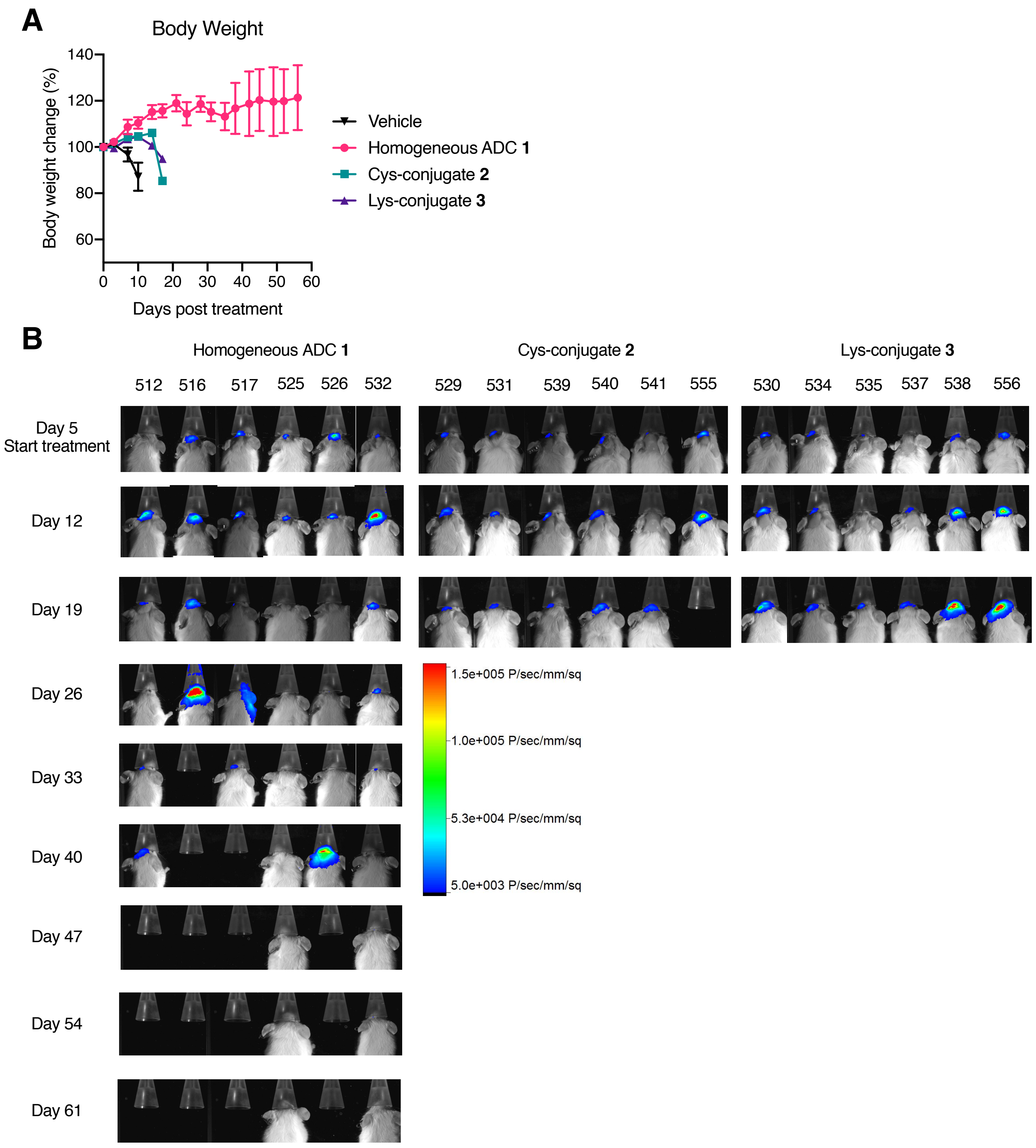


**Figure S3.** Body weight change and bioluminescence imaging in the orthotopic U87ΔEGFR-luc xenograft mouse model. (A) Body weight change during treatment. Vehicle (black inversed triangle), homogeneous ADC **1** (magenta circle), Cys conjugate **2** (green square), and Lys conjugate **3** (purple triangle). Data are presented as mean values ± SEM (n = 6). (B) Bioluminescence images were taken right before ADC administration (Day 5) and then once a week.

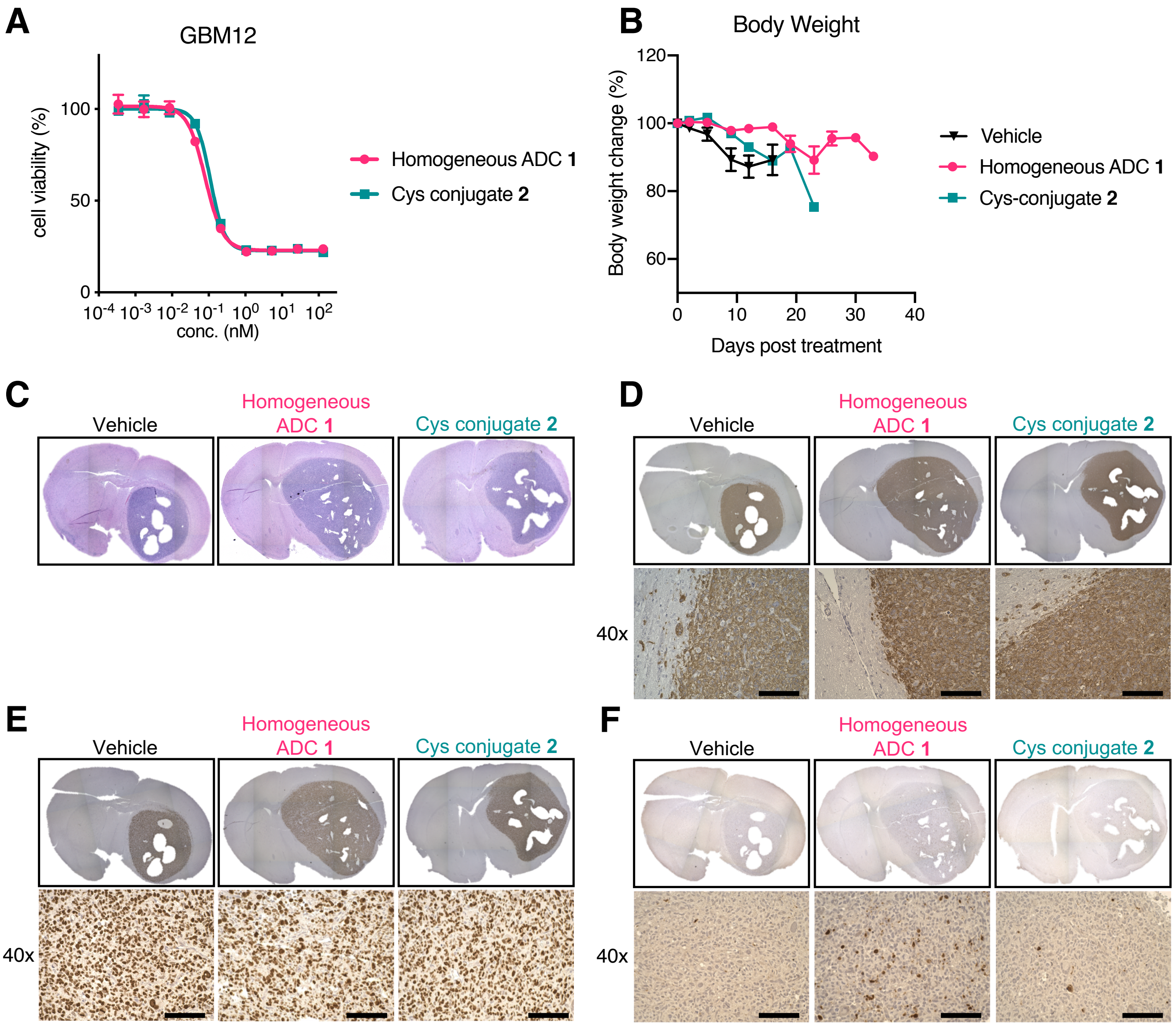


**Figure S4.** In vitro and in vivo evaluation of anti-EGFR ADCs in the GBM12 PDX model. (A) In vitro cell killing potency in GBM12 cells. We tested homogeneous ADC **1** (magenta circle) and Cys conjugate **2** (green square). Concentrations are based on the antibody dose without normalizing to each DAR. All assays were performed in triplicate. (B) Body weight change during treatment in the orthotopic GBM12 xenograft mouse model. Vehicle (n = 15, black inversed triangle), homogeneous ADC **1** (n = 14, magenta circle), and Cys conjugate **2** (n = 14, green square). Data are presented as mean values ± SEM. (C–F) Immunohistochemistry analysis of GBM12 tumors harvested at the terminal stage. Representative images are shown. Tumor sections were stained with (C) H&E, (D) anti-human EGFR, (E) anti-Ki67, and (F) anti-cleaved-caspase 3 antibodies. Scale bar: 100 μm.

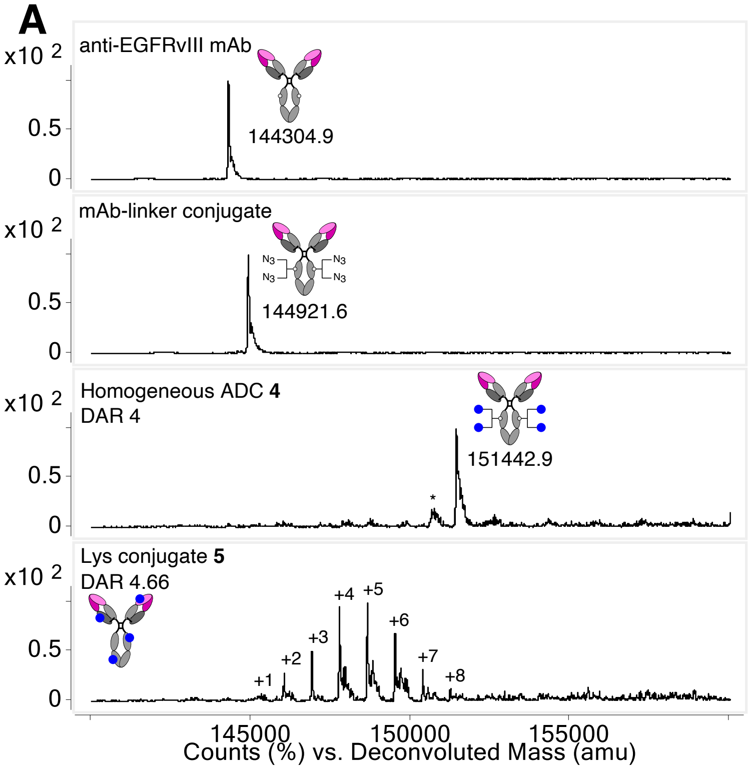


| DAR | 0 | 1 | 2 | 3 | 4 | 5 | 6 | 7 | 8 |
|---|---|---|---|---|---|---|---|---|---|
| Intensity (%) | 0 | 1.3 | 7.1 | 12.9 | 24.4 | 25.4 | 17.8 | 8.1 | 3.0 |

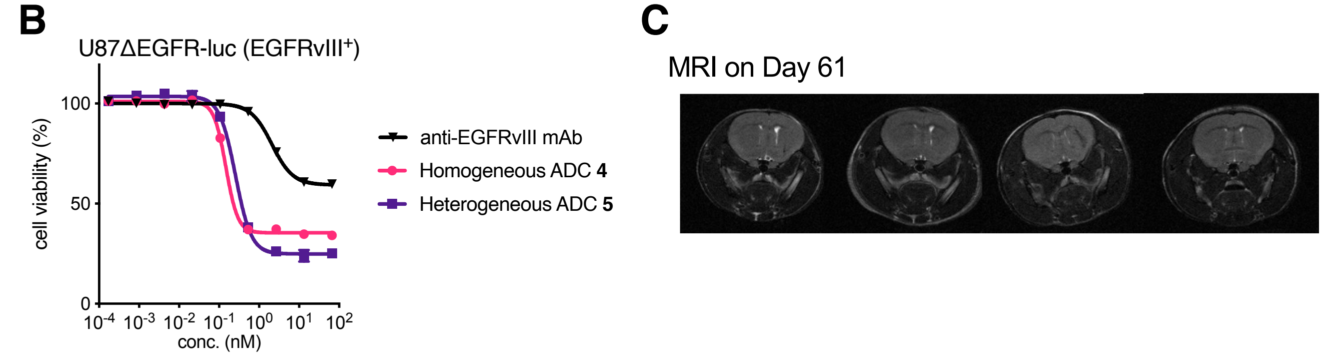


**Figure S5.** Characterization of anti-EGFRvIII ADCs and the anti-tumor effect in vitro and in vivo. (A) Preparation and ESI-MS analysis of homogeneous anti-EGFRvIII ADC **4** and heterogeneous Lys conjugate **5**. First panel: N297A anti-EGFRvIII mAb (depatuxizumab mutant). Second panel: mAb–linker conjugate. Third panel: homogeneous ADC **4** with a DAR of 4. Asterisk (*) indicates a fragment ion detected in ESI-MS analysis. Fourth panel: Lys conjugate **5**. The average DAR was determined to be 4.66 based on the ion intensity of each DAR species. (B) In vitro cell killing potency in U87ΔEGFR-luc cells. We tested unmodified anti-EGFRvIII mAb (black inversed triangle), homogeneous ADC **4** (magenta circle) and heterogeneous ADC **5** (Lys conjugate **5**, purple square). Concentrations are based on the antibody dose without normalizing to each DAR. All assays were performed in triplicate. Data are presented as mean values ± SEM. (C) MRI analysis of 4 survivor mice that were intracranially implanted with U87ΔEGFR-luc cells and treated with a single dose of anti-EGFRvIII ADC **4** at 3 mg/kg on Day 8. The coronal images were taken on Day 61 post tumor implantation. No detectable tumor lesion was observed.

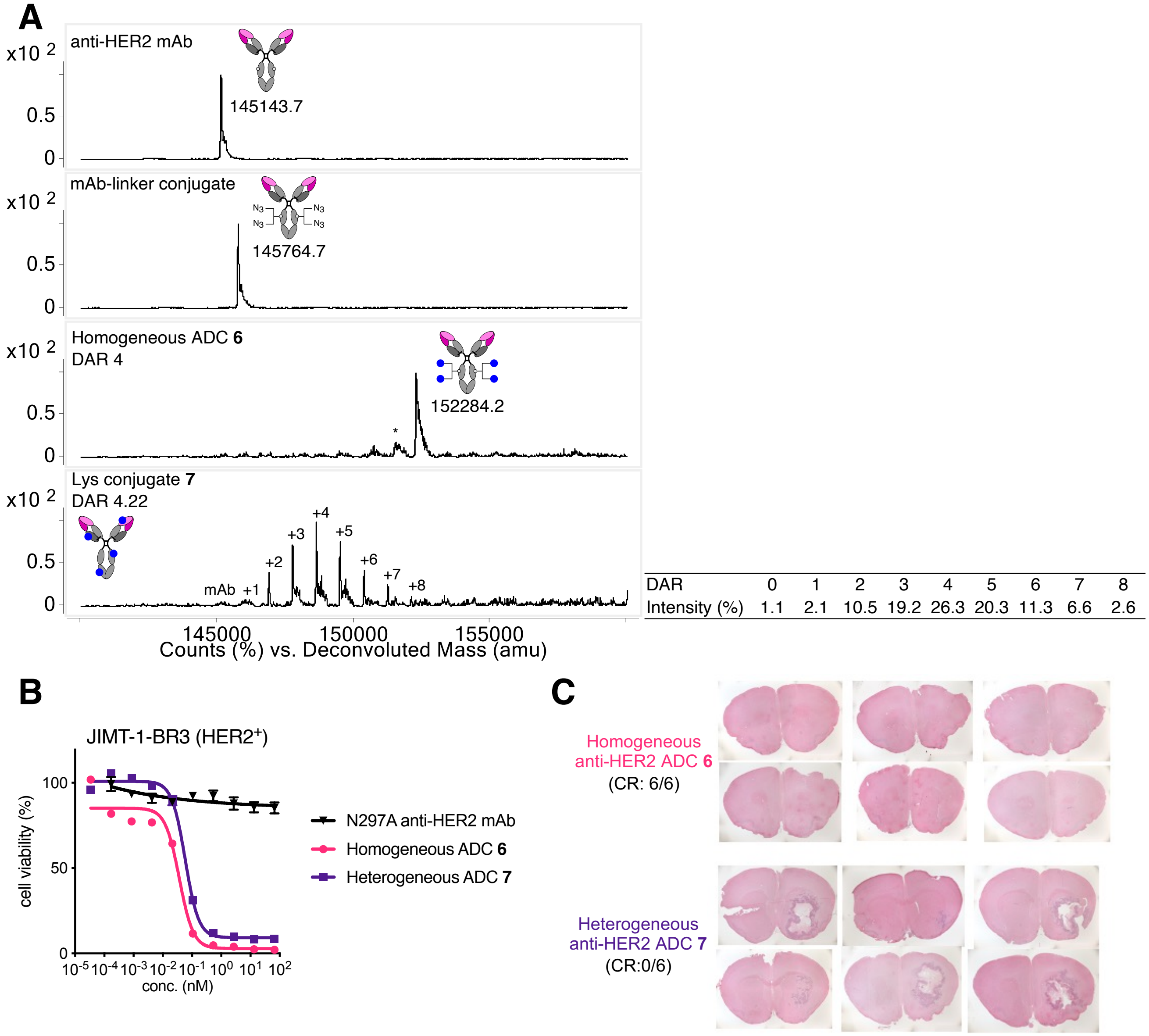


| DAR | 0 | 1 | 2 | 3 | 4 | 5 | 6 | 7 | 8 |
|---|---|---|---|---|---|---|---|---|---|
| Intensity (%) | 1.1 | 2.1 | 10.5 | 19.2 | 26.3 | 20.3 | 11.3 | 6.6 | 2.6 |

**Figure S6.** Characterization of anti-HER2 ADCs and the anti-tumor effect in vitro and in vivo. (A) Preparation and ESI-MS analysis of homogeneous anti-HER2 ADC **6** and heterogeneous Lys conjugate **7**. First panel: N297A anti-HER2 mAb (trastuzumab mutant). Second panel: mAb–linker conjugate. Third panel: homogeneous ADC **6** with a DAR of 4. Asterisk (*) indicates a fragment ion detected in ESI-MS analysis. Fourth panel: Lys conjugate **7**. The average DAR was determined to be 4.22 based on the ion intensity of each DAR species. (B) In vitro cell killing potency in JIMT1-BR3 cells. We tested unmodified anti-HER2 mAb (black inversed triangle), homogeneous ADC **6** (magenta circle) and heterogeneous ADC **7** (Lys conjugate **7**, purple square). Concentrations are based on the antibody dose without normalizing to each DAR. All assays were performed in triplicate. Data are presented as mean values ± SEM. (C) H&E staining of brain tissues. Brain tumor lesions were observed in all mice treated with heterogeneous ADC **7** (6/6) but not in those treated with homogeneous ADC **6** (0/6). CR, complete remission.

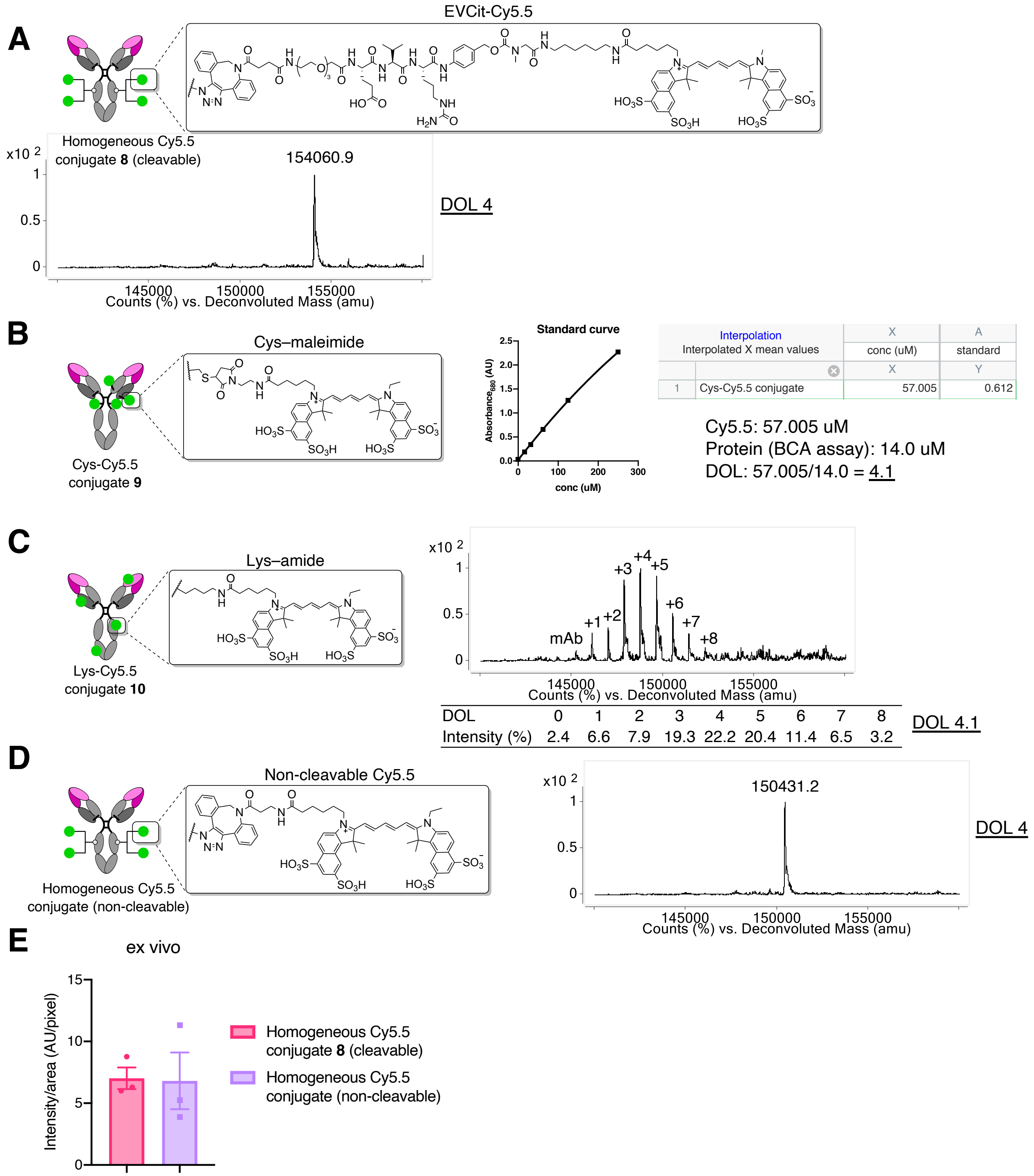


**Figure S7.** Characterization of Cy5.5 conjugates and evaluation for brain tumor targeting. (A) ESI-MS analysis of homogeneous cleavable Cy5.5 conjugate **8** with a DOL of 4. (B) Determination of DOL of Cys-Cy5.5 conjugate **9**. Protein concentration was determined by BCA assay and the molar amount of conjugated Cy5.5 was calculated using a free Cy5.5-based standard curve. The average DOL was then determined to be 4.1 based on the ratio of those values. (C) ESI-MS analysis of Lys-Cy5.5 conjugate **10**. The average DOL was determined to be 4.1 based on the ion intensity of each DOL species. (D) ESI-MS analysis of the homogeneous non-cleavable Cy5.5 conjugate with a DOL of 4. (E) Semi-quantification of the Cy5.5 signal detected in the whole orthotopic U87ΔEGFR-luc brain tumors treated with cleavable conjugate **8** or non-cleavable variant. Data are presented as mean values ± SEM.

**A** DAR 0 mAb-Cy5.5 conjugate **11**

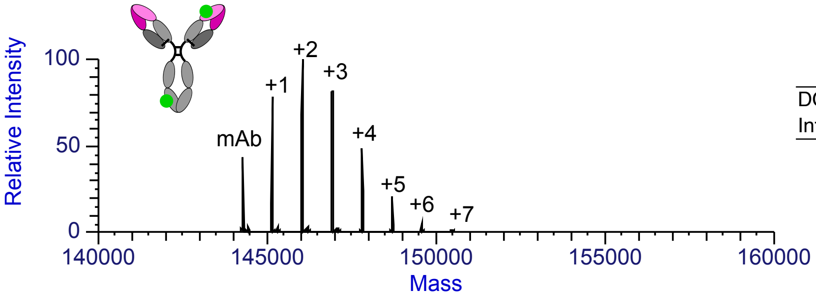


| DOL | 0 | 1 | 2 | 3 | 4 | 5 | 6 | 7 |
|---|---|---|---|---|---|---|---|---|
| Intensity (%) | 11.5 | 20.8 | 26.6 | 21.6 | 12.7 | 5.4 | 1.3 | 0.2 |

→ DOL 2.3

**B** DAR 4 ADC-Cy5.5 conjugate **12**

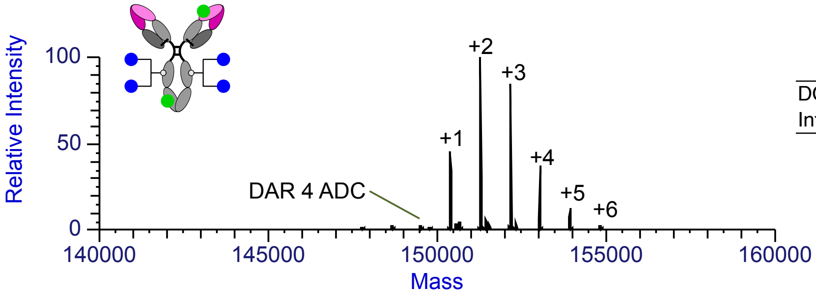


| DOL | 0 | 1 | 2 | 3 | 4 | 5 | 6 |
|---|---|---|---|---|---|---|---|
| Intensity (%) | 0.6 | 16.0 | 35.6 | 29.8 | 13.0 | 4.3 | 0.6 |

→ DOL 2.5

**C** DAR 6 ADC-Cy5.5 conjugate **13**

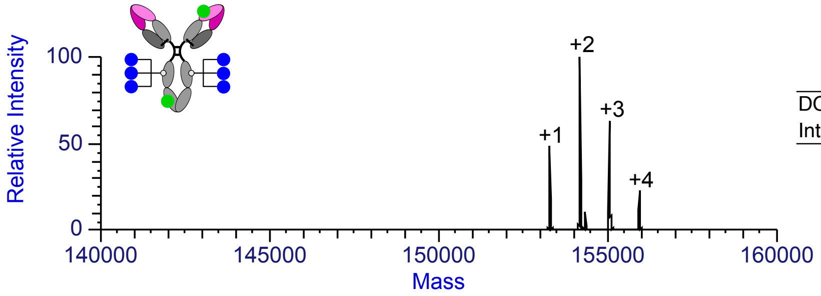


| DOL | 0 | 1 | 2 | 3 | 4 |
|---|---|---|---|---|---|
| Intensity (%) | 0 | 20.9 | 42.7 | 26.7 | 9.7 |

→ DOL 2.3

**D** DAR 8 ADC-Cy5.5 conjugate **14**

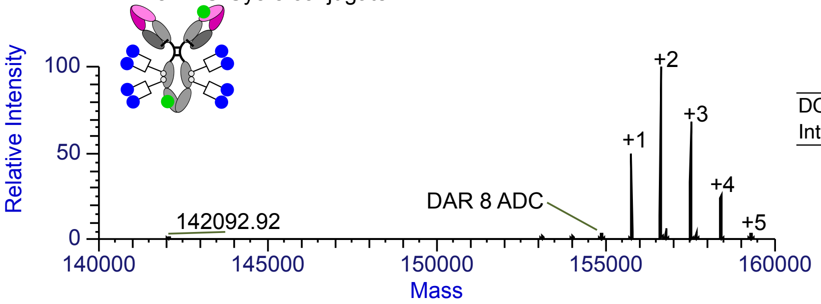


| DOL | 0 | 1 | 2 | 3 | 4 | 5 |
|---|---|---|---|---|---|---|
| Intensity (%) | 1.2 | 19.7 | 40.5 | 27.2 | 10.2 | 1.2 |

→ DOL 2.3

**Figure S8.** Characterization of Cy5.5-labeled MMAF conjugates **11**–**14**. (A) ESI-MS spectra of anti-EGFRvIII mAb-Cy5.5 conjugate **11** with an average DOL of 2.3, (B) DAR 4 MMAF-Cy5.5 conjugate **12** with an average DOL of 2.5, (C) DAR 6 MMAF-Cy5.5 conjugate **13** with an average DOL of 2.3, and (D) DAR 8 MMAF-Cy5.5 conjugate **14** with an average DOL of 2.3.

### Supplementary Tables

**Table S1** $K_D$ values of unmodified mAb and ADCs (n = 3).

| | $K_D$ (nM) | CI95 (nM) |
|---|---|---|
| anti-EGFR mAb (cetuximab mutant) | 0.039 | 0.036–0.043 |
| Homogeneous ADC **1** | 0.047 | 0.042–0.053 |
| Cys conjugate **2** | 0.044 | 0.040–0.048 |
| Lys conjugate **3** | 0.045 | 0.038–0.052 |

Calculated based on the data shown in Figure 2A.

**Table S2** $EC_{50}$ values of ADCs in GBM cell lines (n = 3).

| | $EC_{50}$ (nM) | |
|---|---|---|
| | U87ΔEGFR | Gli36δEGFR |
| anti-EGFR mAb (cetuximab mutant) | 1.99 (1.54–2.65) | – |
| Homogeneous ADC **1** | 0.072 (0.064–0.083) | 0.048 (0.044–0.052) |
| Cys conjugate **2** | 0.110 (0.100–0.122) | 0.035 (0.033–0.037) |
| Lys conjugate **3** | 0.140 (0.126–0.156) | 0.048 (0.044–0.053) |

Calculated based on the data shown in Figure 2B. Values in parentheses are 95% confidential intervals.

**Table S3** Summary of *in vivo* PK (n = 3).

| | $t_{1/2\beta\ \mathrm{Total\ mAb}}$ (day) | $t_{1/2\beta\ \mathrm{ADC}}$ (day) | $AUC_{0\text{-}14\ \mathrm{Total\ mAb}}$ (µg/mL×day) | $AUC_{0\text{-}14\ \mathrm{ADC}}$ (µg/mL×day) | $CL_{\mathrm{obs\ Total\ mAb}}$ (mg/kg)/(µg/mL)/day | $CL_{\mathrm{obs\ ADC}}$ (mg/kg)/(µg/mL)/day |
|---|---|---|---|---|---|---|
| anti-EGFR mAb (cetuximab mutant) | 10.9 | – | 146.1 (127.5 – 164.7) | – | 0.0119 | – |
| Homogeneous ADC **1** | 9.8 | 8.6 | 140.7 (129.2 – 152.1) | 145.2 (135.0 – 155.4) | 0.0126 | 0.0133 |
| Cys conjugate **2** | 7.8 | 4.2 | 129.6 (121.8 – 137.4) | 97.2 (91.0 – 103.4) | 0.0167 | 0.0279 |
| Lys conjugate **3** | 10.4 | 8.9 | 129.0 (117.5 – 140.5) | 150.0 (141.2 – 158.9) | 0.0137 | 0.0132 |

Calculated based on the data shown in Fig. 5A and 5B. Values in parentheses are 95% confidential intervals. AUC, area under the curve; CL, clearance.

**Table S4** Statistical significance.

| Main Figures | Method | Asterisk | Comparison | *P* value |
|---|---|---|---|---|
| **Figure 3B** | Log-rank[a] (Mantel-Cox) | ** | Vehicle vs Homogeneous ADC **1** | *P* = 0.0032 |
| | | ** | Vehicle vs Cys conjugate **2** | *P* = 0.0032 |
| | | ** | Vehicle vs Lys conjugate **3** | *P* = 0.0032 |
| | | ** | Homogeneous ADC **1** vs Cys conjugate **2** | *P* = 0.0046 |
| | | ** | Homogeneous ADC **1** vs Lys conjugate **3** | *P* = 0.0046 |
| | | ns | Cys conjugate **2** vs Lys conjugate **3** | *P* = 0.9000 |
| **Figure 3D** | Log-rank[a] (Mantel-Cox) | **** | Vehicle vs Homogeneous ADC **1** | $P = 6.8 \times 10^{-8}$ |
| | | ns | Vehicle vs Cys conjugate **2** | *P* = 0.12 |
| | | **** | Homogeneous ADC **1** vs Cys conjugate **2** | $P = 1.017 \times 10^{-5}$ |
| **Figure 3F** | Tukey–Kramer | **** | Vehicle vs Homogeneous ADC **1** | *P* < 0.0001 |
| | | ** | Vehicle vs Cys conjugate **2** | *P* = 0.0021 |
| | | * | Homogeneous ADC **1** vs Cys conjugate **2** | *P* = 0.0279 |
| **Figure 3G** | Tukey–Kramer | * | Vehicle vs Homogeneous ADC **1** | *P* = 0.0149 |
| | | * | Vehicle vs Cys conjugate **2** | *P* = 0.0452 |
| | | ns | Homogeneous ADC **1** vs Cys conjugate **2** | *P* = 0.8010 |
| **Figure 3H** | Tukey–Kramer | ** | Vehicle vs Homogeneous ADC **1** | *P* = 0.0064 |
| | | ns | Vehicle vs Cys conjugate **2** | *P* = 0.6793 |
| | | * | Homogeneous ADC **1** vs Cys conjugate **2** | *P* = 0.0333 |
| **Figure 4B** | Log-rank (Mantel-Cox) | * | Homogeneous ADC **4** vs Lys conjugate **5** | *P* = 0.0195 |
| **Figure 5D** | Tukey–Kramer | ** | Homogeneous Cy5.5 conjugate **8** vs Cys-Cy5.5 conjugate **9** | *P* = 0.0045 |
| | | ** | Homogeneous Cy5.5 conjugate **8** vs Lys-Cy5.5 conjugate **10** | *P* = 0.0020 |
| | | ns | Cys-Cy5.5 conjugate **9** vs Lys-Cy5.5 conjugate **10** | *P* = 0.6589 |
| **Figure 5F Kidney** | Tukey–Kramer | * | Homogeneous Cy5.5 conjugate **8** vs Cys-Cy5.5 conjugate **9** | *P* = 0.0144 |
| | | ns | Homogeneous Cy5.5 conjugate **8** vs Lys-Cy5.5 conjugate **10** | *P* = 0.9037 |
| | | * | Cys-Cy5.5 conjugate **9** vs Lys-Cy5.5 conjugate **10** | *P* = 0.0235 |
| **Figure 5G Liver** | Tukey–Kramer | * | Homogeneous Cy5.5 conjugate **8** vs Cys-Cy5.5 conjugate **9** | *P* = 0.0357 |
| | | ns | Homogeneous Cy5.5 conjugate **8** vs Lys-Cy5.5 conjugate **10** | *P* = 0.8889 |
| | | ns | Cys-Cy5.5 conjugate **9** vs Lys-Cy5.5 conjugate **10** | *P* = 0.0632 |
| **Figure 6E** | Dunnett's test | * | Day +1 Homogeneous Cy5.5 conjugate **8** vs Cys-Cy5.5 conjugate **9** | *P* = 0.0153 |
| | | * | Day +1 Homogeneous Cy5.5 conjugate **8** vs Lys-Cy5.5 conjugate **10** | *P* = 0.0267 |
| | | * | Day +3 Homogeneous Cy5.5 conjugate **8** vs Cys-Cy5.5 conjugate **9** | *P* = 0.0188 |
| | | * | Day +3 Homogeneous Cy5.5 conjugate **8** vs Lys-Cy5.5 conjugate **10** | *P* = 0.0202 |
| | | * | Day +5 Homogeneous Cy5.5 conjugate **8** vs Cys-Cy5.5 conjugate **9** | *P* = 0.0318 |
| | | ns | Day +5 Homogeneous Cy5.5 conjugate **8** vs Lys-Cy5.5 conjugate **10** | *P* = 0.3210 |
| **Figure 7C** | Dunnett's test | ns | DAR 4 ADC **12** vs DAR 0 mAb **11** | *P* = 0.8483 |
| | | * | DAR 4 ADC **12** vs DAR 6 ADC **13** | *P* = 0.0396 |
| | | * | DAR 4 ADC **12** vs DAR 8 ADC **14** | *P* = 0.0288 |
| **Figure 7E** | Dunnett's test | *** | DAR 4 ADC **12** vs DAR 0 mAb **11** | *P* = 0.0006 |
| | | ns | DAR 4 ADC **12** vs DAR 6 ADC **13** | *P* = 0.2450 |
| | | ns | DAR 4 ADC **12** vs DAR 8 ADC **14** | *P* = 0.7237 |

[a] *P* values were adjusted by the Bonferroni correction for multiple comparisons. *$P < 0.05$; **$P < 0.01$; ***$P < 0.001$; ****$P < 0.0001$.

### Supplementary Notes

#### General information

Unless otherwise noted, all materials for chemical synthesis were purchased from commercial suppliers (Acros Organics, AnaSpec, Broadpharm, Chem-Impex International, Fisher Scientific, Levena Biopharma, Sigma Aldrich, TCI America, and other vendors) and used as received. All anhydrous solvents were purchased and stored over activated molecular sieves under argon atmosphere.

Analytical reverse-phase high performance liquid chromatography (RP-HPLC) was performed using an Agilent LC-MS system consisting of a 1100 HPLC and a 1946D single quadrupole electrospray ionization (ESI) mass spectrometer equipped with a C18 reverse-phase column (Accucore™ C18 column, 3 × 50 mm, 2.6 µm, Thermo Scientific) or a Thermo LC-MS system consisting of a Vanquish UHPLC and a LTQ XL™ linear ion trap mass spectrometer equipped with a C18 reverse-phase column (Accucore™ Vanquish™ C18+ UHPLC column, 2.1 × 50 mm, 1.5 µm, Thermo Scientific). Standard analysis conditions for organic molecules were as follows: flow rate = 0.5 mL/min (for both systems); solvent A = water containing 0.1% formic acid; solvent B = acetonitrile containing 0.1% formic acid. Compounds were analyzed using a linear gradient and monitored with UV detection at 210 and 254 nm. Preparative HPLC was performed using a Breeze HPLC system (Waters) equipped with a C18 reverse-phase column (XBridge Peptide BEH C18 OBD Prep Column, 130Å, 5 µm, 19 × 150 mm, Waters). Standard purification conditions were as follows: flow rate = 20 mL/min; solvent A = water containing 0.05% trifluoroacetic acid (TFA), 0.1% formic acid or 0.1% $NH_4OH$; solvent B = acetonitrile containing 0.05% TFA (standard conditions), 0.1% formic acid (FA conditions), or 0.1% $NH_4OH$ (basic conditions). Compounds were analyzed using a linear gradient and monitored with UV detection at 210 and 254 nm. In all cases, fractions were analyzed off-line using either of the LC-MS systems for purity confirmation and those containing a desired product were lyophilized using a Labconco Freezone 4.5 Liter Benchtop Freeze Dry System. High-resolution mass spectra were obtained using an Agilent 6530 Accurate Mass Q-TOF LC/MS or a Thermo Q Exactive™ Hybrid Quadrupole-Orbitrap™ Mass Spectrometer.

### Synthesis

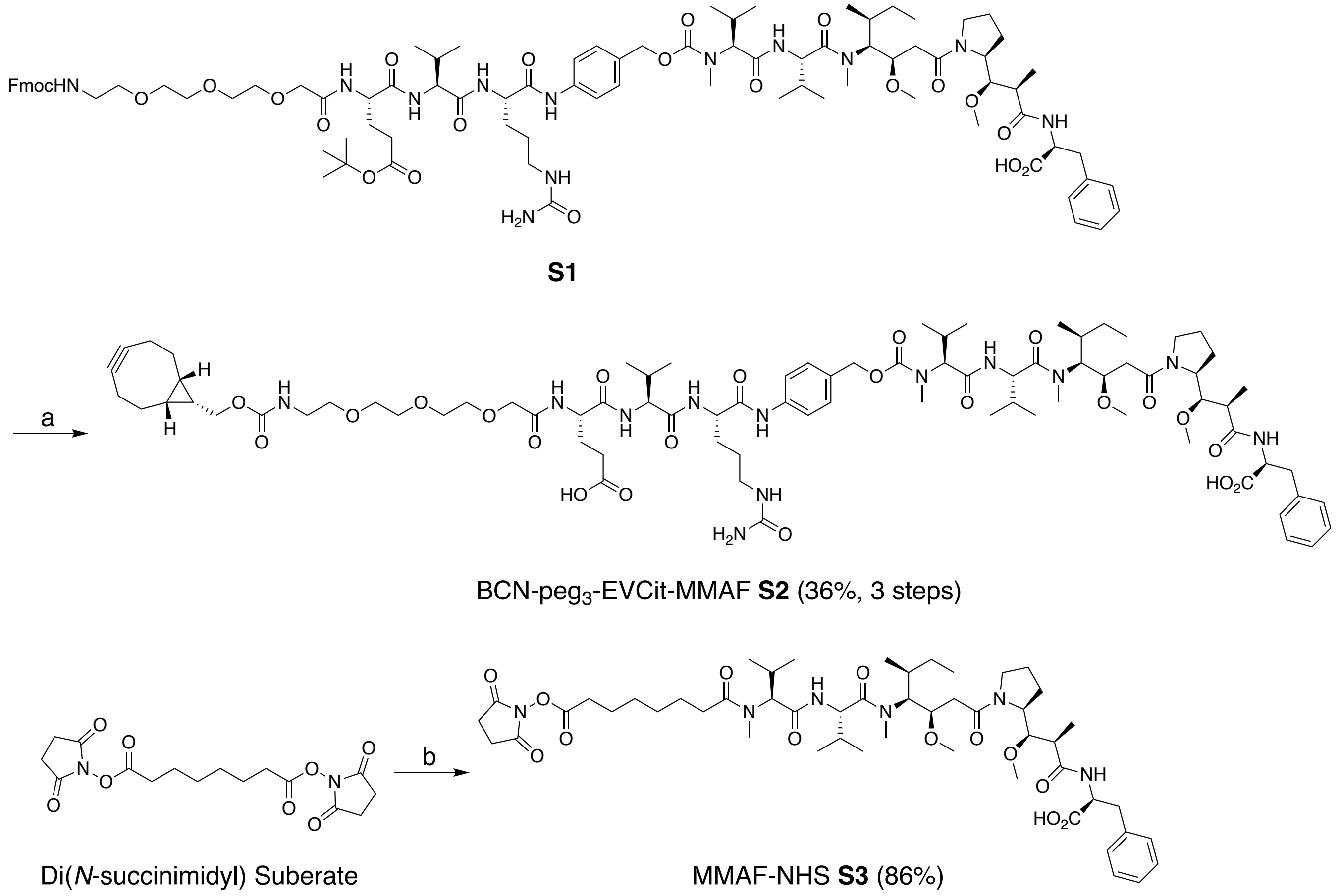


Synthesis of BCN-EVCit-MMAF module **S2** and MMAF-NHS ester (**S3**). Reagents and conditions: (a) 50% diethylamine/DMF, rt, 1 h, and 50% TFA/DCM, rt, 1 h, then MMAF, DIPEA, HOAt, DMF, rt, overnight. (b) MMAF, DIPEA, HOAt, di(*N*-succinimidyl) suberate, 37 ℃, overnight.

### Preparation of BCN-peg$_3$-EVCit-PABC-MMAF (S2)

Fmoc-peg$_3$-E(O*t*-Bu)VCit-PABC-MMAF (**S1**, 15 mg, 8.7 μmol, prepared as described previously (Yamazaki et al., 2021)) was dissolved in 50% diethylamine/DMF solution at room temperature. After 1 h, the solution was concentrated in vacuo and used in the next step without further purification. The crude products were dissolved in 50% TFA/DCM solution at room temperature. After being stirred at room temperature for 1 h, the solution was concentrated in vacuo and the crude compounds were precipitated with cold diethyl ether (5–6 mL) followed by centrifugation at 2,000 × *g* for 3 min (3 times). BCN-NHS (3.8 mg, 13.1 μmol, Berry&Associates) and DIPEA (7.6 μL, 43.5 μmol) were added to a solution of this crude mixture in DMF (1 mL) and the mixture was stirred at room temperature overnight. The crude products were purified by preparative RP-HPLC under basic conditions to afford analytically pure peptide **S2** (5.1 mg, 36% for the 3 steps). Purity was confirmed by LC-MS. White powder. HRMS (ESI) Calcd. For $C_{82}H_{126}N_{12}O_{22}Na_2$ $[M+2Na]^{2+}$: 838.4447, Found: 838.4467.

**Preparation of MMAF-NHS ester (S3)**

DIPEA (42.5 µL, 0.24 mmol) was added to a solution of MMAF (102.9 mg, 0.12 mmol), di(*N*-succinimidyl) suberate (224.5 mg, 0.61 mmol), 1-hydroxy-7-azabenzotriazole (HOAt, 16.6 mg, 0.12 mmol) in DMF (2 mL) and the mixture was stirred at 37 ºC overnight. The crude products were purified by preparative RP-HPLC to afford analytically pure peptide **S3** (103.7 mg, 86%). Purity was confirmed by LC-MS. White powder. HRMS (ESI) Calcd. For $C_{51}H_{80}N_6O_{13}Na$ $[M+Na]^+$: 1007.5676, Found: 1007.5676.

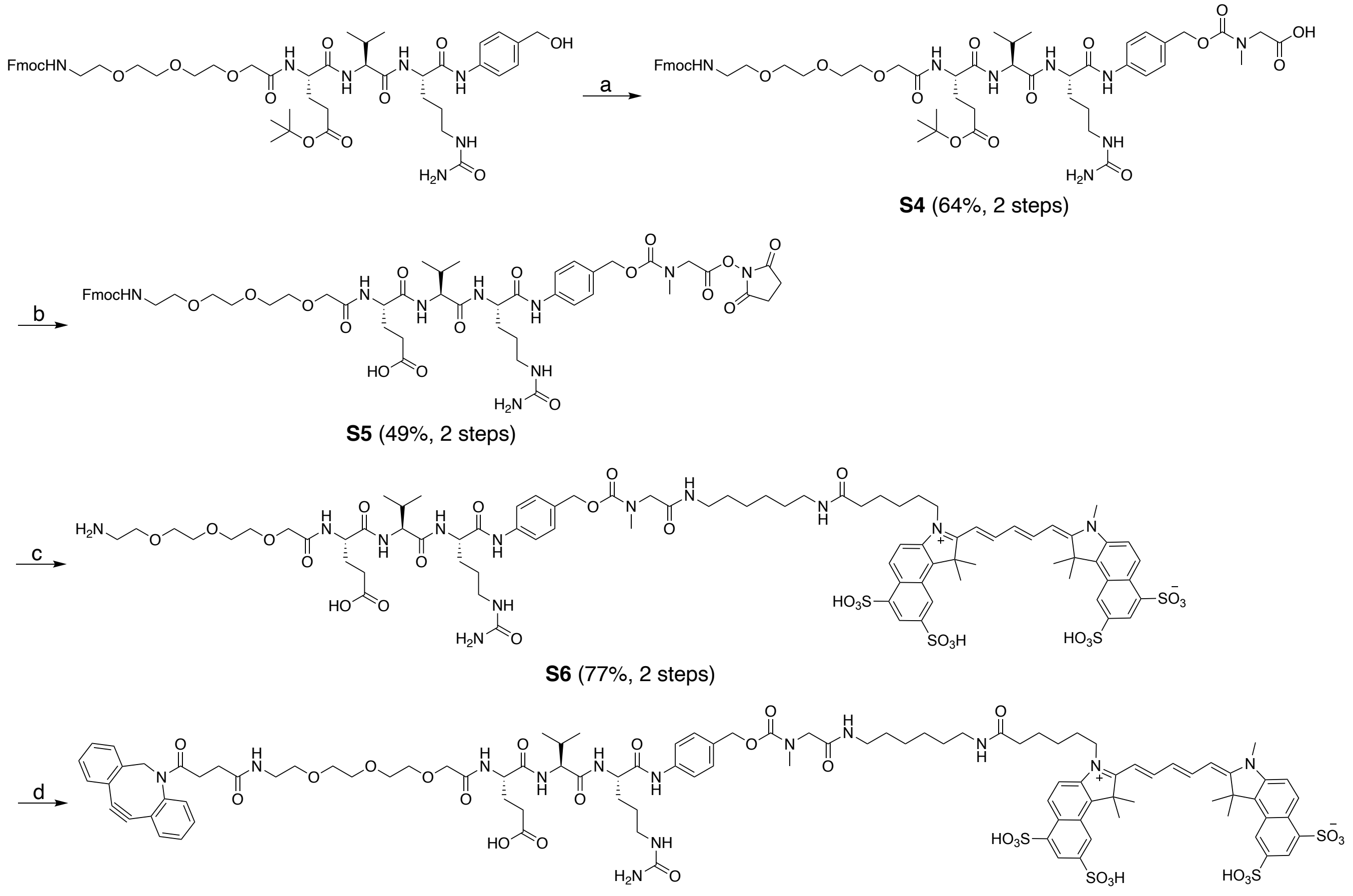


Synthesis of DBCO-EVCit-Cy5.5 module **S7**. Reagents and conditions: (a) bisPNP, DMAP, DMF, rt, 2 h, then sarcosine, rt, 1h. (b) NHS, EDC·HCl, DCM, then 10% TFA/DCM. (c) Cy5.5-amine, DIPEA, rt, overnight, then diethylamine. (d) DBCO-NHS, DIPEA, DMF, DMSO, rt, overnight.

**Preparation of Fmoc-peg$_3$-E(O*t*-Bu)VCit-PABC-sarcosine (S4)**

Bis(2,4-dinitrophenyl) carbonate (52.9 mg, 174 µmol) and DMAP (8.5 mg, 69.6 µmol) were added to a solution of Fmoc-peg$_3$-E(O*t*-Bu)VCit-PABOH (34 mg, 34.8 µmol) in DMF (1 mL), and the mixture was stirred at room temperature for 2 h. To the mixture were added a solution of sarcosine (93 mg, 1.0 mmol) in water (1 mL) and additional DMF (0.5 mL) and the mixture was stirred at room temperature for 1 h. The crude products were purified by preparative RP-HPLC to afford analytically pure peptide **S4** (24.2 mg, 64%). Purity was confirmed by LC-MS. White powder. HRMS (ESI) Calcd. For $C_{54}H_{74}N_8O_{16}Na$ $[M+Na]^+$: 1113.5115, Found: 1113.5111.

**Preparation of Fmoc-peg$_3$-EVCit-PABC-sarcosine NHS ester (S5)**

Fmoc-peg$_3$-E(O*t*-Bu)VCit-PABC-sarcosine (**S4**, 24.2 mg, 22.2 μmol), NHS (7.7 mg, 66.6 μmol), and EDC·HCl (12.8 mg, 66.6 μmol) were dissolved in DCM (1.5 mL) and the mixture was stirred at room temperature for 4 h. Then the reaction mixture was quenched with 15% citric acid and extracted with DCM. The organic layer was washed with brine, dried over $Na_2SO_4$, and concentrated. The crude products were dried in vacuo and used immediately in the next step without purification. The crude products were dissolved in 10% TFA/DCM solution. After being stirred at room temperature for 30 min, the solution was concentrated in vacuo and the residue was purified by preparative RP-HPLC to afford analytically pure peptide **S5** (12.4 mg, 49% for the 2 steps). Purity was confirmed by LC-MS. White powder. HRMS (ESI) Calcd. For $C_{54}H_{70}N_9O_{18}$ $[M+H]^+$: 1132.4833, Found: 1132.4826.

**Preparation of H$_2$N-peg$_3$-EVCit-PABC-sarcosine-Cy5.5 (S6)**

A solution of Cy5.5-amine (500 μL, 10 mM in DMSO, 5 μmol) and DIPEA (3.5 μL, 20 μmol) was added to a solution of Fmoc-peg$_3$-EVCit-PABC-sarcosine NHS ester (**S5**, 500 μL, 10 mM in DMSO, 5 μmol) and the mixture was stirred at room temperature for 1.5 h. Additional NHS ester **S5** (200 μL, 10 mM in DMSO, 2 μmol) was added and the mixture was stirred at room temperature overnight. Diethylamine (800 μL) was added and the reaction mixture was stirred at room temperature for 1 h. The solution was concentrated in vacuo and the residue was purified by preparative RP-HPLC to afford analytically pure peptide **S6** (6.9 mg, 77% for the 2 steps). Purity was confirmed by LC-MS. Blue powder. HRMS (ESI) Calcd. For $C_{81}H_{108}N_{12}O_{26}S_4$ $[M–2H]^{2–}$: 896.3196, Found: 896.3188.

**Preparation of DBCO-peg$_3$-EVCit-PABC-sarcosine-Cy5.5 (S7)**

H$_2$N-peg$_3$-EVCit-PABC-sarcosine-Cy5.5 (**S6**, 6.2 mg, 3.45 μmol) was dissolved in DMF (500 μL) and DMSO (300 μL). To the solution were added DIPEA (1.2 μL, 6.9 μmol) and DBCO-NHS ester (1.8 mg, 4.49 μmol), and the mixture was stirred in the dark at room temperature overnight. The crude products were purified by preparative RP-HPLC under basic conditions to afford analytically pure peptide **S7** (6.5

mg, 90%). Purity was confirmed by LC-MS. Blue powder. HRMS (ESI) Calcd. For $C_{100}H_{120}N_{13}O_{28}S_4$ $[M–3H]^{3–}$: 692.9088, Found: 692.9085.

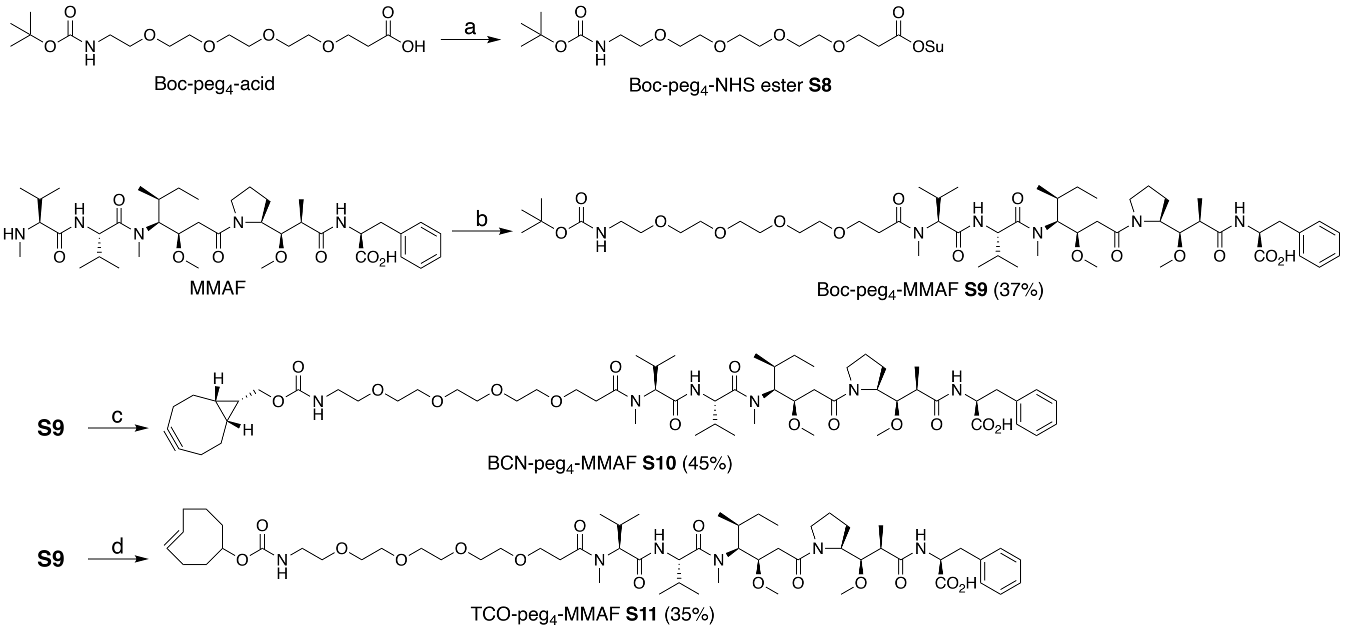


Synthesis of non-cleavable MMAF modules **S10** and **S11**. Reagents and conditions: (a) NHS, EDC·HCl, DCM, rt, overnight. (b) **S8**, HOAt, DIPEA, DMF, 37 ºC, overnight. (c) 50% TFA/DCM, rt, 30 min, then BCN-NHS, DIPEA, DMF, rt, 3 h. (d) 50% TFA/DCM, rt, 30 min, then TCO-NHS, DIPEA, DMF, rt, 3 h.

**Synthesis of Boc-peg$_4$-MMAF (S9)**

Boc-peg$_4$-acid (38.1 mg, 104 µmol), NHS (23.9 mg, 208 µmol), and EDC·HCl (39.7 mg, 208 µmol) were dissolved in DCM (0.75 mL) and the mixture was stirred at room temperature overnight. Then the reaction mixture was quenched with 15% citric acid and extracted with DCM. The organic layer was washed with brine, dried over $Na_2SO_4$, and concentrated. Crude Boc-peg4-NHS ester (**S8**) were dried in vacuo and used immediately in the next step without purification.

DIPEA (10 µL, 57.4 µmol) was added to a solution of MMAF (24.3 mg, 28.7 µmol), crude NHS ester **S8** (20 mg, 43 µmol, 100 mg/mL solution in DMSO), and HOAt (7.8 mg, 57.4 µmol) in DMF (200 µL) and the mixture was stirred at 37 ºC overnight. The crude products were purified by preparative RP-HPLC to afford analytically pure peptide **S9** (11.5 mg, 37%). Purity was confirmed by LC-MS. White powder. HRMS (ESI) Calcd. For $C_{55}H_{95}N_6O_{15}$ $[M+H]^+$: 1079.6850, Found: 1079.6829.

**Synthesis of BCN-peg$_4$-MMAF (S10)**

Boc-peg$_4$-MMAF (**S9**, 11.5 mg, 10.7 µmol) was dissolved in 50% TFA/DCM solution. After being stirred at room temperature for 30 min, the solution was concentrated in vacuo and the crude compounds were precipitated with cold diethyl ether (10 mL) followed by centrifugation at 2,000 × *g* for 3 min (3 times). The residue was equally aliquoted into two vials for BCN and TCO installation, respectively.

For BCN installation, BCN-NHS (2.0 mg, 6.96 µmol, Berry&Associates) and DIPEA (0.9 µL, 10.7 µmol) were added to a solution of this crude mixture in DMF (300 µL) and the mixture was stirred at room temperature for 3 h. The crude products were purified by preparative RP-HPLC under FA conditions to afford analytically pure peptide **S10** (2.8 mg, 45% for the 2 steps). Purity was confirmed by LC-MS. White powder. HRMS (ESI) Calcd. For $C_{61}H_{99}N_6O_{15}$ $[M+H]^+$: 1155.7163, Found: 1155.7148. TCO-peg$_4$-MMAF (**S11**) was synthesized in a similar manner.

**TCO-peg$_4$-MMAF (S11)**

TCO-NHS (1.9 mg, 6.96 µmol) was used instead of BCN-NHS. 2.1 mg, 35% yield for the 2 steps. Purity was confirmed by LC-MS. White powder. HRMS (ESI) Calcd. For $C_{59}H_{99}N_6O_{15}$ $[M+H]^+$: 1131.7163, Found: 1131.7151.

### LC-MS data

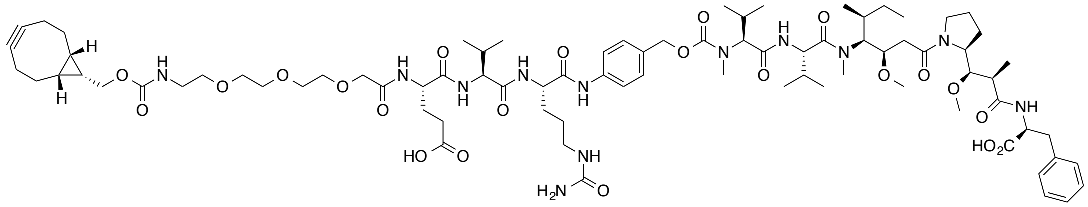


BCN-peg$_3$-EVCit-PABC-MMAF (**S2**)

Print of all graphic windows

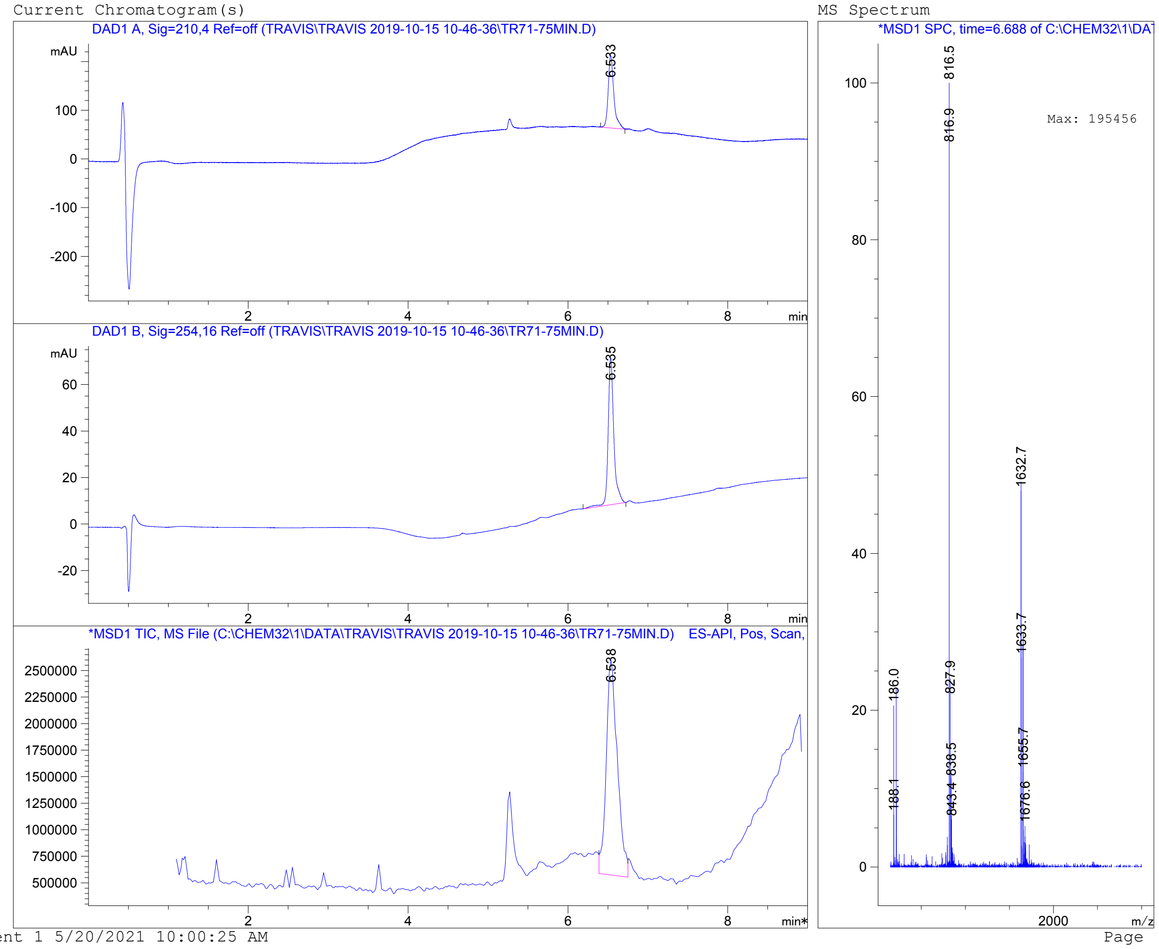


Instrument 1 5/20/2021 10:00:25 AM Page 1 of 1

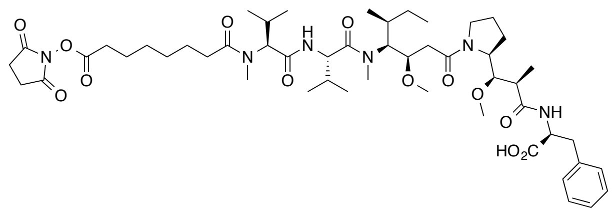


MMAF-NHS ester (**S3**)

Print of all graphic windows

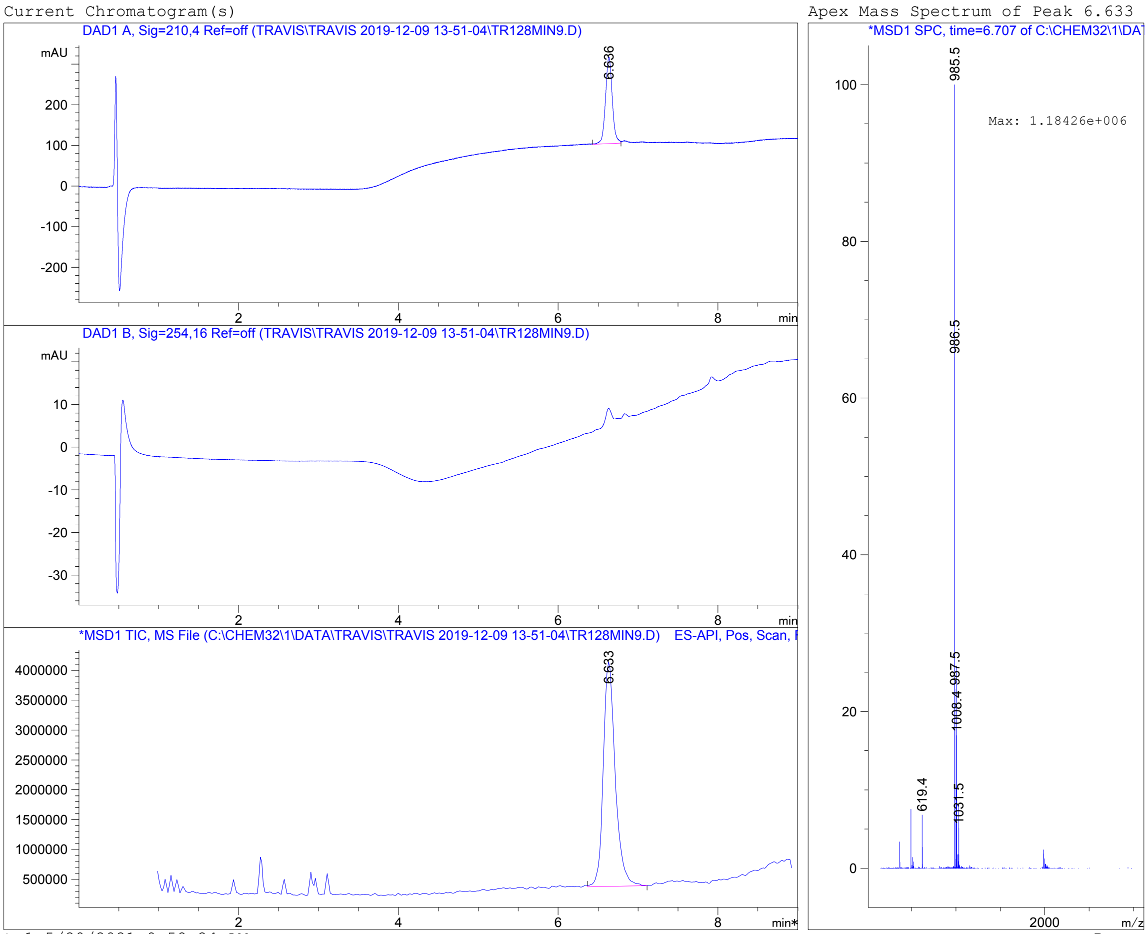


Instrument 1 5/20/2021 9:58:24 AM

Page 1 of 1

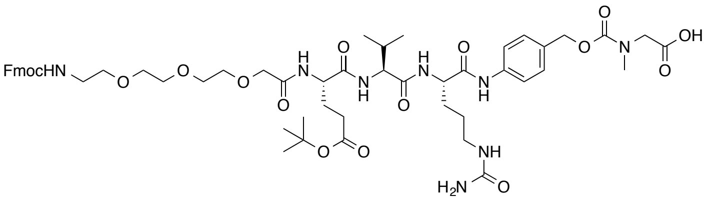

Fmoc-peg$_3$-E(O*t*-Bu)VCit-PABC-sarcosine (**S4**)

Print of all graphic windows

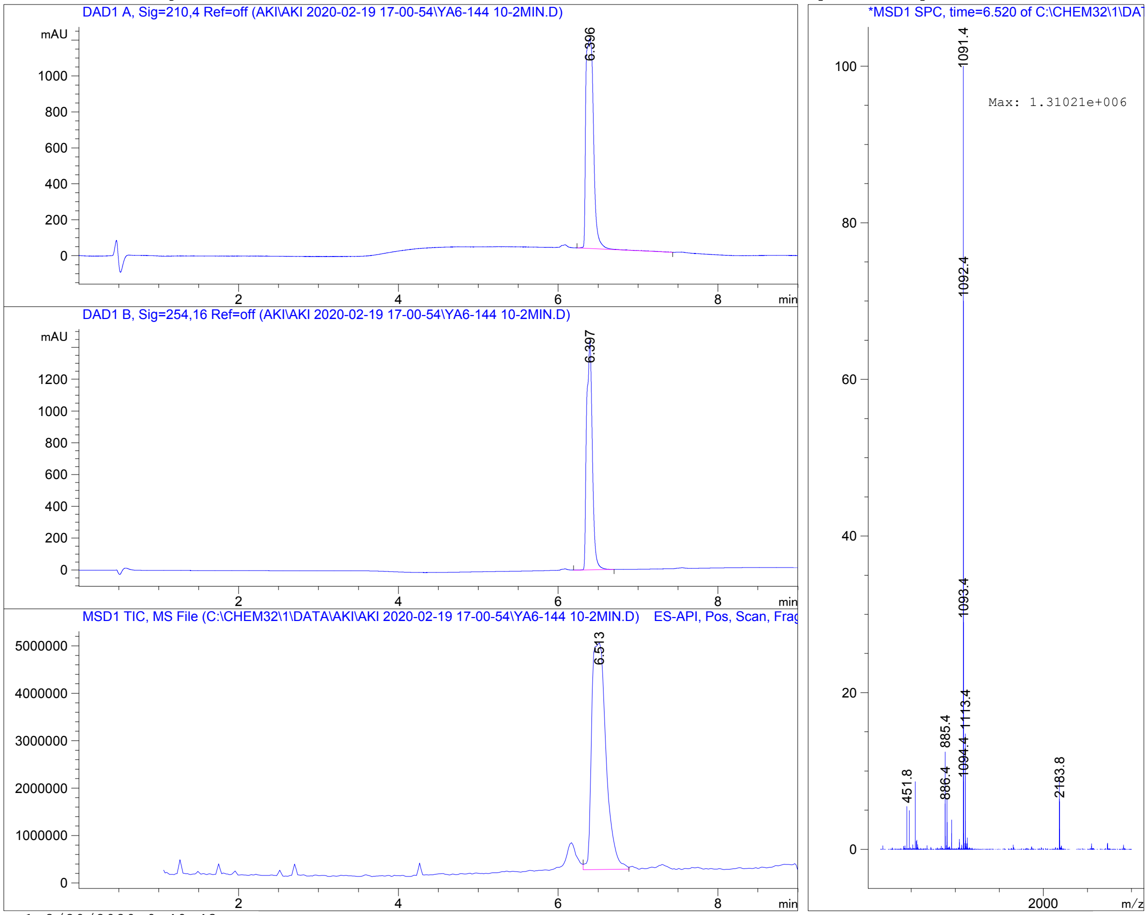


Instrument 1 2/20/2020 9:49:43 AM Page 1 of 1

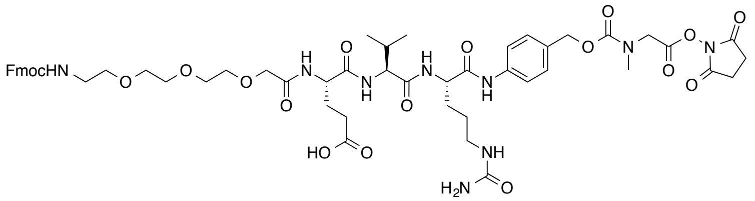

Fmoc-peg$_3$-EVCit-PABC-sarcosine NHS ester (**S5**)

Print of all graphic windows

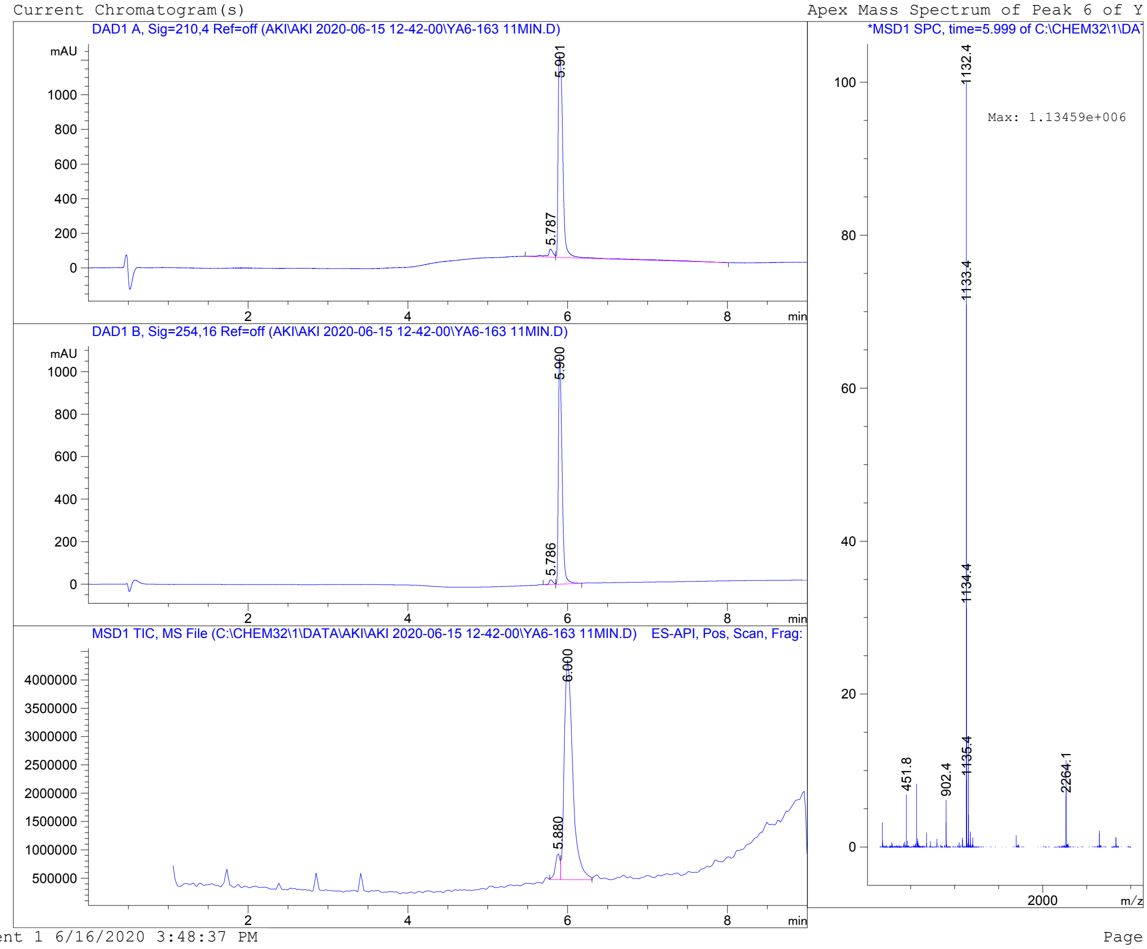


Instrument 1 6/16/2020 3:48:37 PM

Page 1 of 1

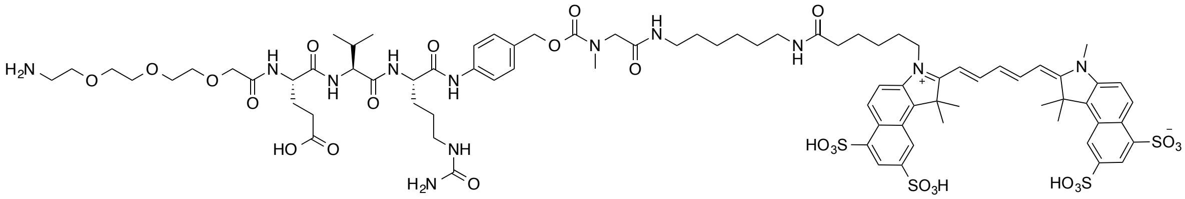


$H_2N$-peg$_3$-EVCit-PABC-sarcosine-Cy5.5 (**S6**)

Print of all graphic windows

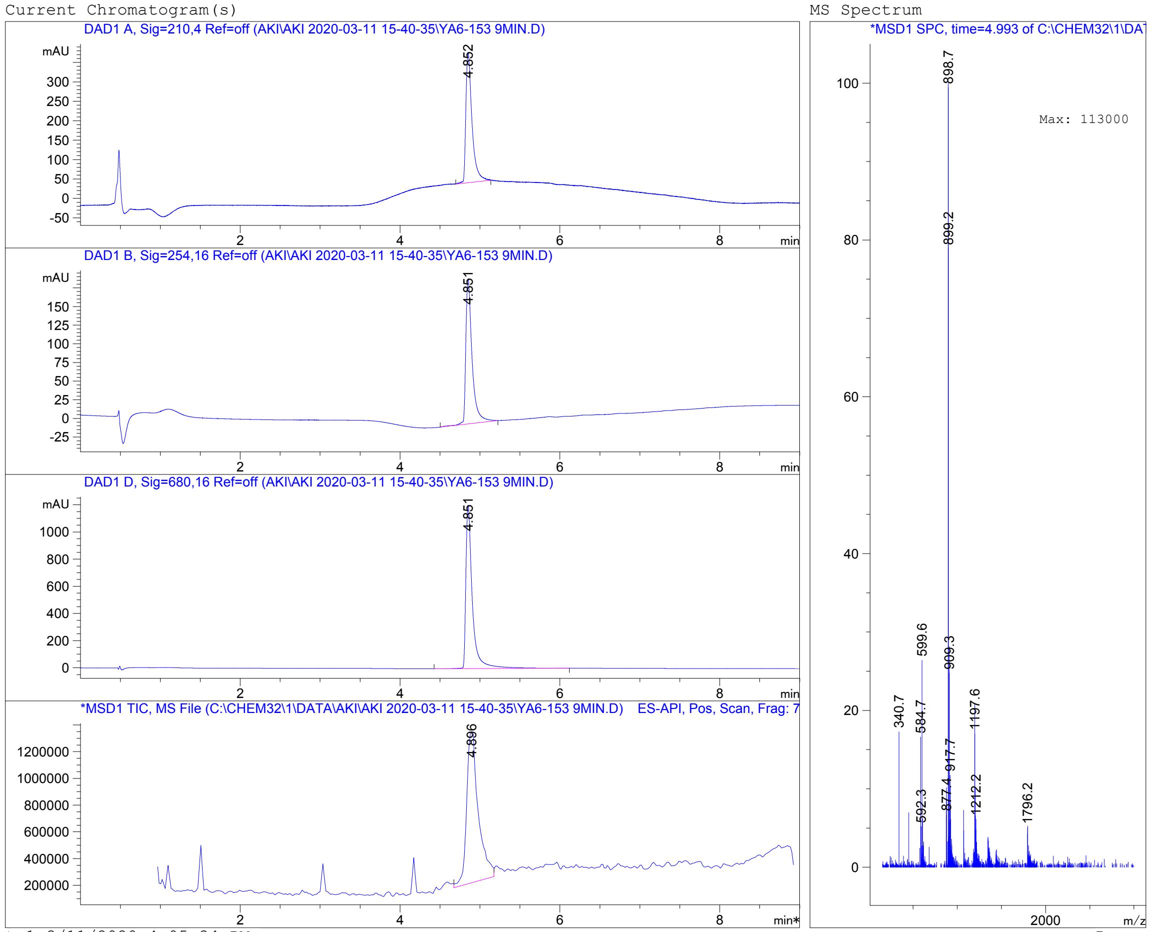


Instrument 1 3/11/2020 4:05:34 PM    Page 1 of 1

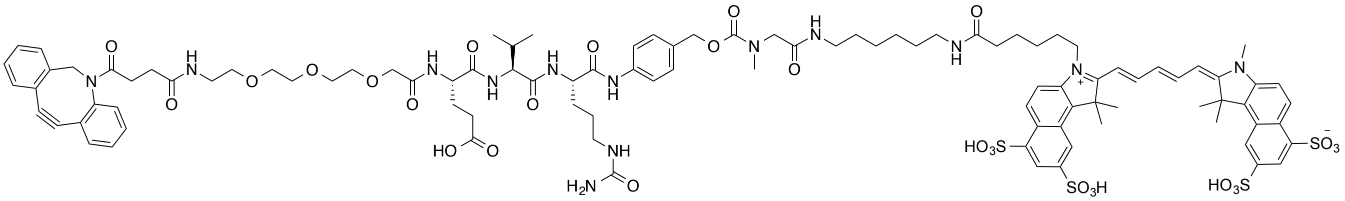

DBCO-peg$_3$-EVCit-PABC-sarcosine-Cy5.5 (**S7**)

Print of all graphic windows

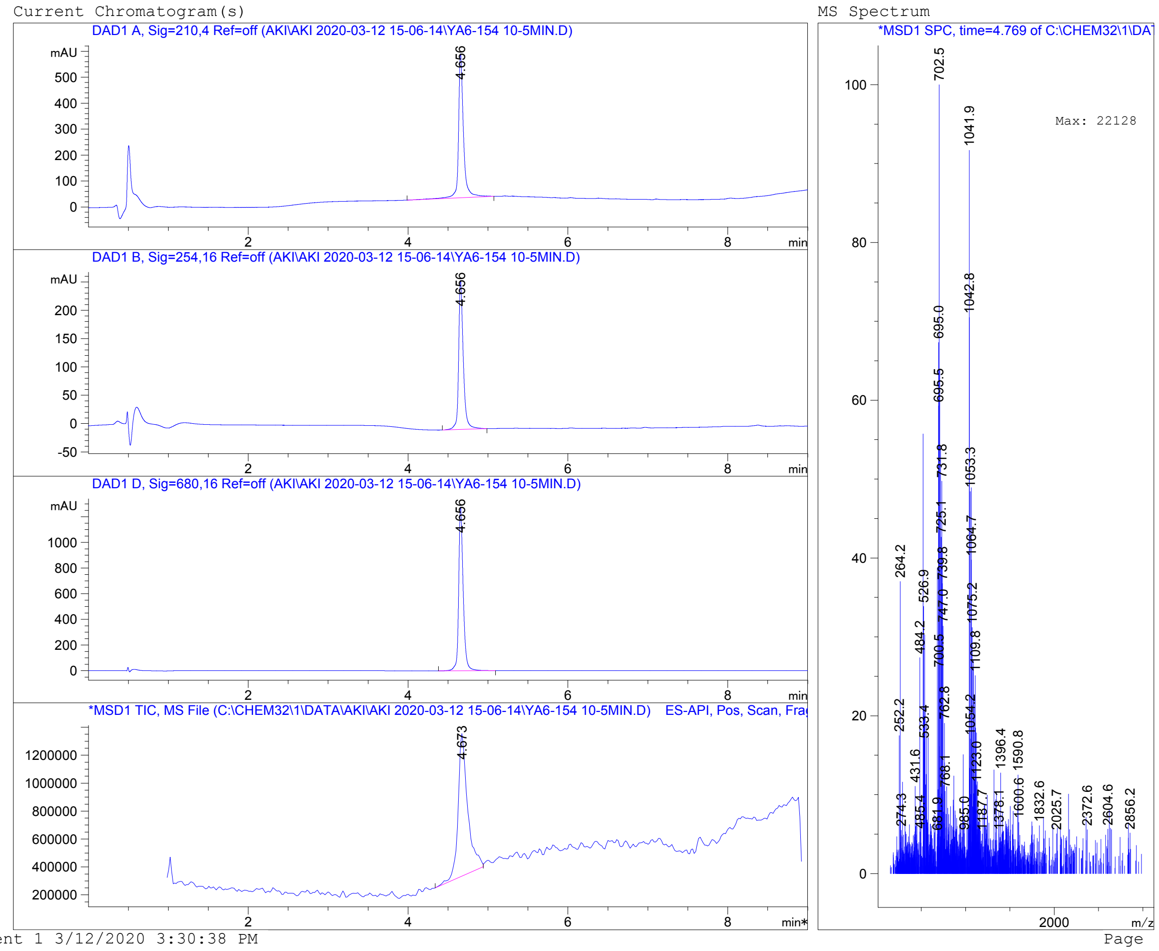


Instrument 1 3/12/2020 3:30:38 PM Page 1 of 1

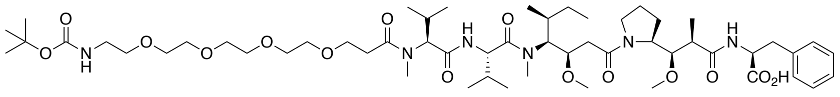

Boc-peg$_4$-MMAF (**S9**)

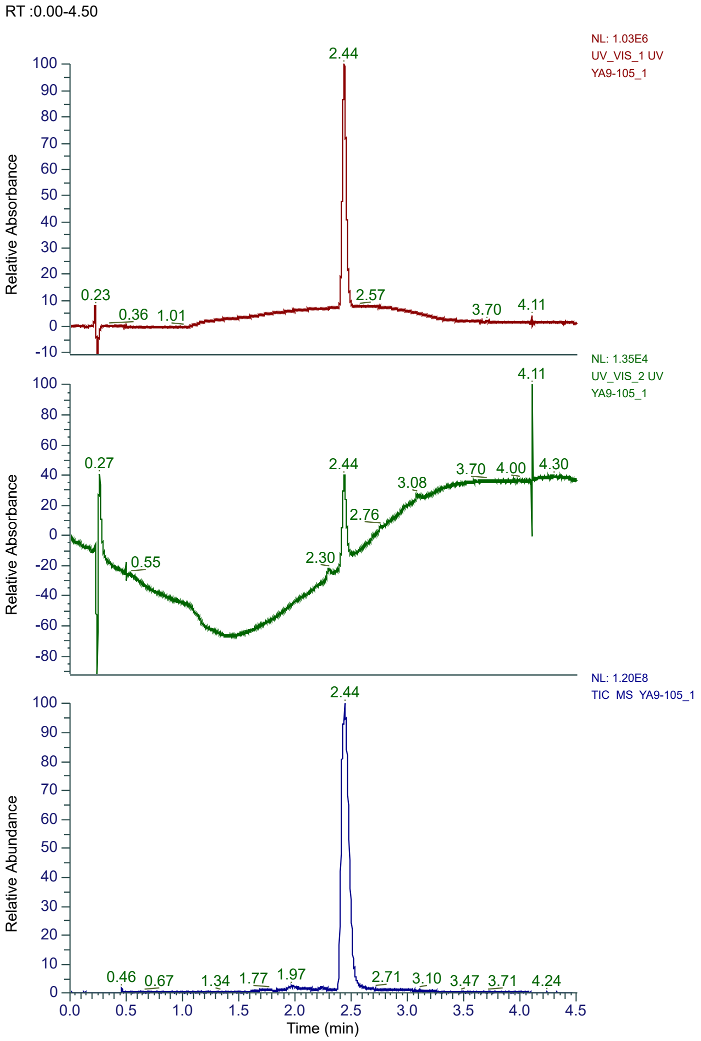


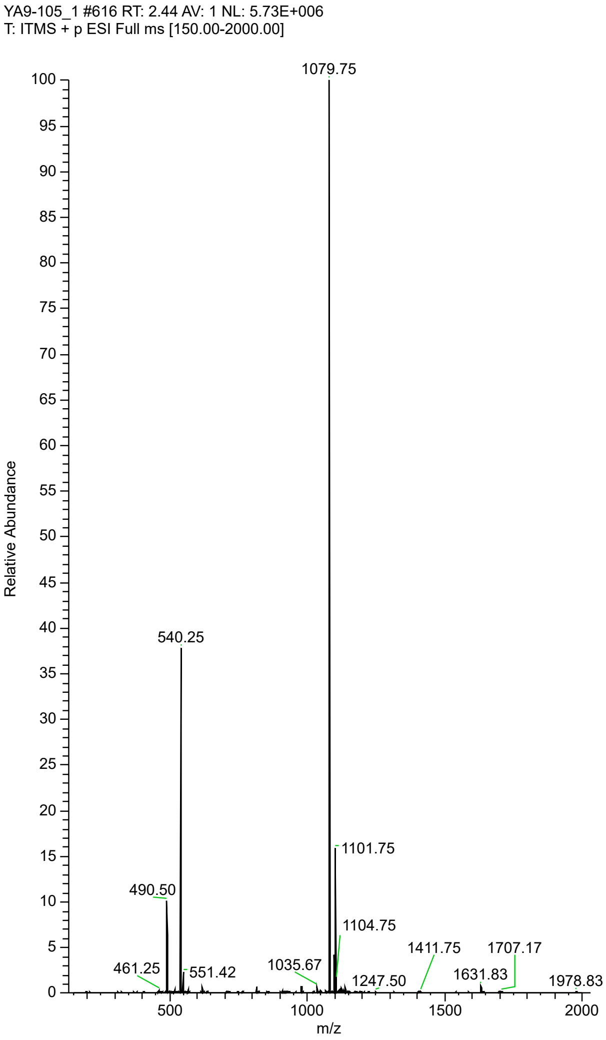

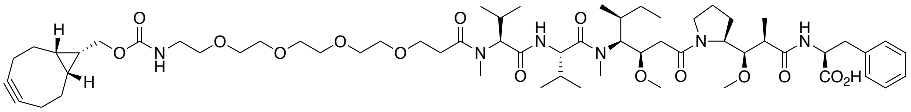

BCN-peg$_4$-MMAF (**S10**)

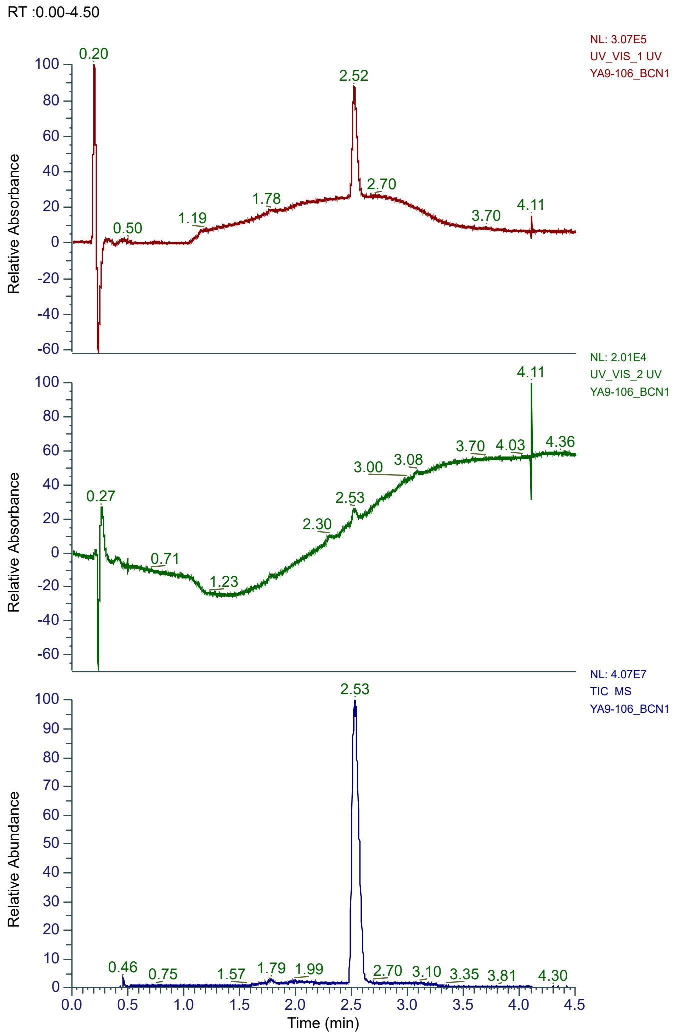


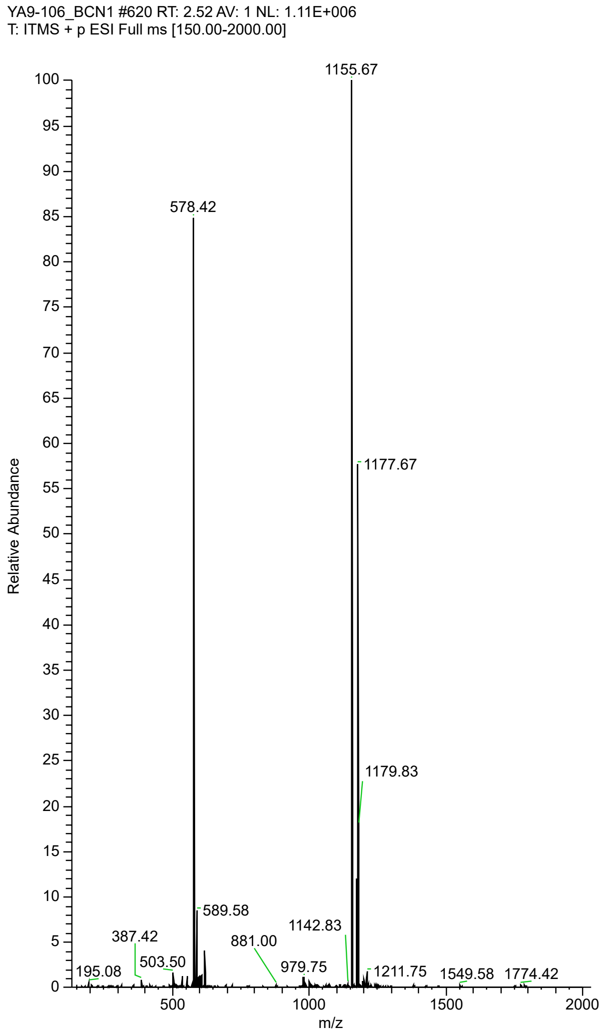

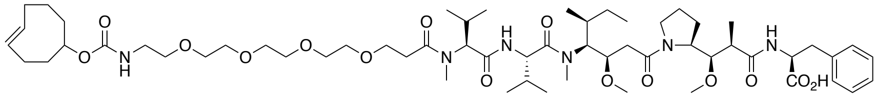

TCO-peg$_4$-MMAF (**S11**)

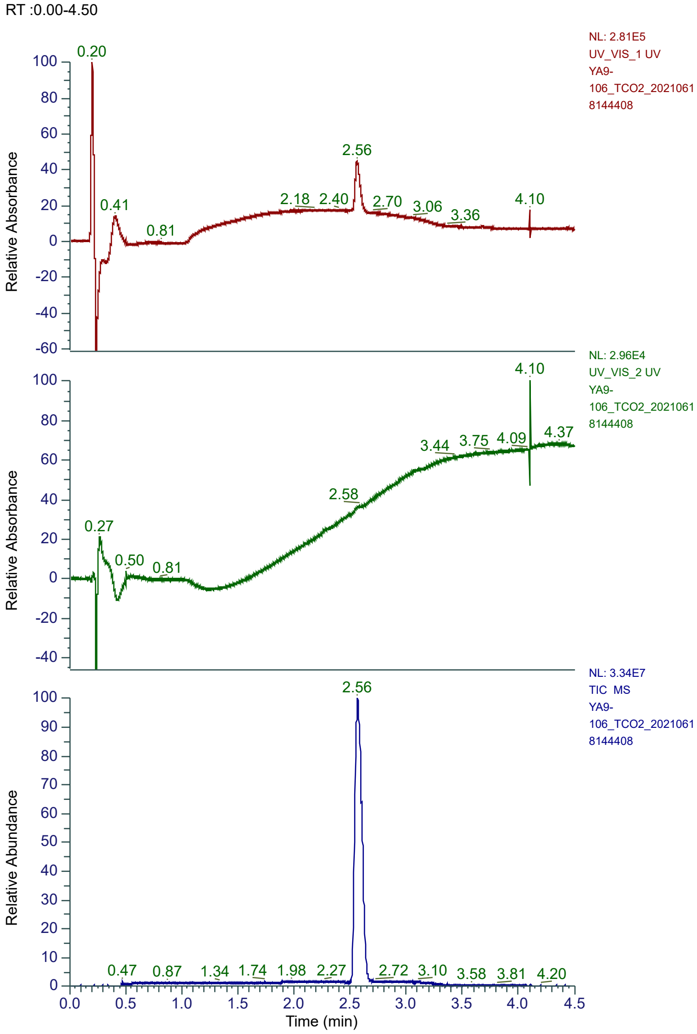


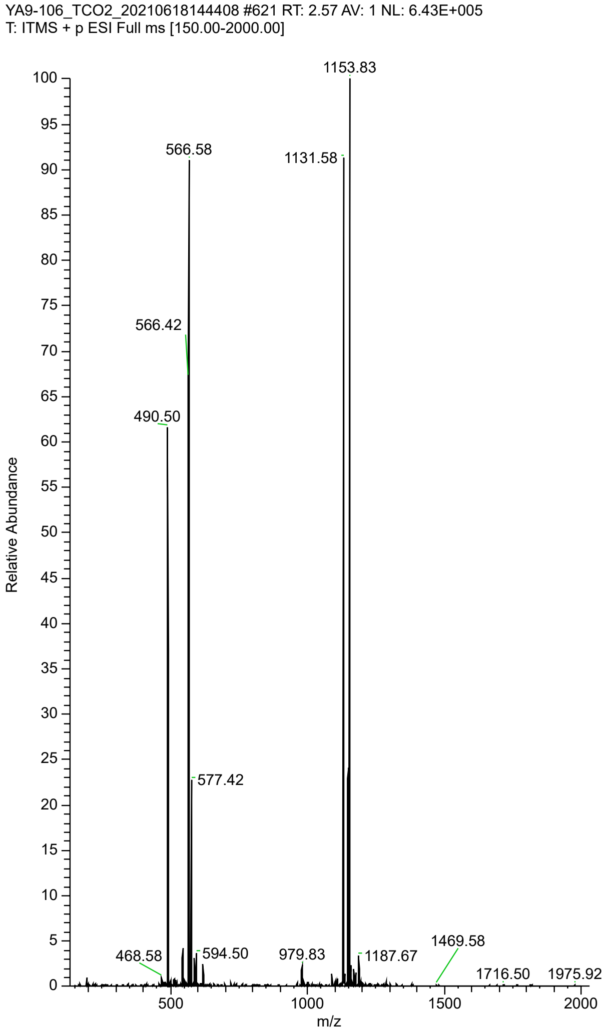
